# *Escherichia coli* metabolism under short-term repetitive substrate dynamics: Adaptation and trade-offs

**DOI:** 10.1101/2020.03.14.982140

**Authors:** Eleni Vasilakou, Mark C. M. van Loosdrecht, S. Aljoscha Wahl

## Abstract

**Background:** Microbial metabolism is highly dependent on the environmental conditions. Especially, the substrate concentration, as well as oxygen availability, determine the metabolic rates. In large-scale bioreactors, microorganisms encounter dynamic conditions in substrate and oxygen availability (mixing limitations), which influence their metabolism and subsequently their physiology. Earlier, single substrate pulse experiments were not able to explain the observed physiological changes generated under large-scale industrial fermentation conditions.

**Results:** In this study we applied a repetitive feast-famine regime in an aerobic *Escherichia coli* culture in a time-scale of seconds. The regime was applied for several generations, allowing cells to adapt to the (repetitive) dynamic environment. The observed response was highly reproducible over the cycles, indicating that cells were indeed fully adapted to the regime. We observed an increase of the specific substrate and oxygen consumption (average) rates during the feast-famine regime, compared to a steady-state (chemostat) reference environment. The increased rates at same (average) growth rate led to a reduced biomass yield (30% lower). Interestingly, this drop was not followed by increased by-product formation, pointing to the existence of energy-spilling reactions and/or less effective ATP synthesis. During the feast-famine cycle, the cells rapidly increased their uptake rate. Within 10 seconds after the beginning of the feeding, the substrate uptake rate was higher (4.68 μmol/g_CDW_/s) than reported during batch growth (3.3 μmol/g_CDW_/s). The high uptake led to an accumulation of several intracellular metabolites, during the feast phase, accounting for up to 34 % of the carbon supplied. Although the metabolite concentrations changed rapidly, the cellular energy charge remained unaffected, suggesting well-controlled balance between ATP producing and ATP consuming reactions. The role of inorganic polyphosphate as an energy buffer is discussed.

**Conclusions:** The adaptation of the physiology and metabolism of *Escherichia coli* under substrate dynamics, representative for large-scale fermenters, revealed the existence of several cellular mechanisms coping with stress. Changes in the substrate uptake system, storage potential and energy-spilling processes resulted to be of great importance. These metabolic strategies consist a meaningful step to further tackle reduced microbial performance, observed under large-scale cultivations.

## 1. Introduction

Microorganisms are widely used for the production of chemicals, ranging from small organic acids to large proteins, including biopharmaceuticals, biochemicals and bulk biofuels [1–3]. In order to meet the cost targets and demands, large-scale production cultivations are and will be required [4]. However, scale-up of microbial processes is not a trivial process, as strain performance usually declines from lab to industrial-scale biorectors [5–7]. One root of this problem is the mixing limitations, which characterize large-scale bioreactors, and lead to several heterogeneities in the cultivation environment. Important parameters, affected by the constraints in mass and heat transfer, are nutrient concentrations, pH, dissolved gases, temperature and other parameters, which have been extensively reported in many studies and reviews [8–13].

Substrate gradients are frequently considered as a main reason for performance reduction. Commonly, the substrate concentration should be kept at low levels to avoid overflow metabolism [14–17]. To achieve such conditions, fed-batch or chemostat regimes are applied. At large scale with mixing limitations, this leads to varying concentration of the substrate in different areas of the reactor [8, 18–20]. Especially, for most large-scale bioreactors, there is one feeding inlet; close to that point the substrate concentration is (very) high, while it becomes lower in the other parts of the reactor. Thus, cells inside the reactor circulate between zones of substrate excess and zones of substrate limitation. Depending on the scale and the type of the reactor, the timeframes are in order of seconds to minutes [11, 12, 18]. The presence and origin of these gradients has also been demonstrated by computational fluid dynamic simulations [18, 21–23].

It is known that such heterogeneities have a big impact on the cellular metabolism, from physiology to metabolic fluxes and gene expression, and subsequently on product formation. Cells traveling close to the feeding zone will increase their substrate uptake rates, which may lead to increased overflow metabolism leading to unwanted products and energy spilling. Additionally, the high uptake rate leads to high oxygen demands, potentially inducing oxygen depletion in this zone [24–26]. On the other hand, cells passing through areas with low substrate concentration may re-consume overflow metabolites and/or activate stress reponse pathways, due to substrate limitations [12]. For example, *Escherichia coli* cultures, cyclically circulating from high to low glucose levels, have shown decreased biomass yields and by-product formation [8, 19].

Two approaches to improve the cell performance in large-scale bioprocesses are: 1) Optimization of bioreactor design and operating conditions preventing gradients, and/or 2) Development of more robust strains which can cope with these conditions.

Especially, for designing more robust strains a mechanistic understanding of the (metabolic) responses to the environment dynamics is required. The cellular regulation mechanisms to cope with frequently changing environments have been, and still are, key questions in microbiology, not only for industrial applications, but also for understanding microbial ecosystems such as the natural habitat of *E.coli,* involving the lower intestine of humans and animals, water, sediment and soil [27].

The cellular behaviour, under substrate dynamic conditions, has been studied using numerous scale-down experiments, for many types of microorganisms (for reviews check [28, 29]). Commonly these studies derived observations on the physiology, such as average rates of growth, substrate uptake, product formation and respiration, but focused less on the metabolic network and the underlying mechanisms of the cellular responses. For example, the energetic state of the cell, storage accummulation, futile cycles and more phenomena, occuring under dynamic conditions, need further investigation. In addition, the fact that microorganisms, cultivated in large-scale bioreactors, face substrate gradients in a cyclic mode (alternating from substrate excess to limitation), has been highly neglected. Most of the scale-down experiments assessed the behaviour of the culture shortly after applying a single perturbation event. However, we strongly consider that cells develop different adaptation strategies, when facing variations in environmental conditions, over long time periods (not long enough for genetic evolution). Several researchers studied such conditions [30–34] and suggested significant changes in physiology, metabolic fluxes, as well as transcription and translation, when the cells moved between different stress zones repeatedly, eventually leading to reduced growth and productivity [6]. Therefore, the long-term responses to successive substrate gradients will be different than those which would occur in short-term (< 5 generations) or after a sudden perturbation. Suarez-Mendez CA*, et al.* [35] applied repetitive glucose perturbations to *Saccharomyces cerevisiae* culture, using a feast-famine regime, proving that the dynamic responses of the adapted culture showed many differences compared to stimulus-response experiments, such as the absence of the ATP paradox [36]. A similar study has been performed for *Penicillium chrysogenum* [37]. However, only a few studies have been previously performed with *E.coli*, assessing the effects of substrate gradients in long-term. Pickett AM*, et al.* [38] were the first ones to apply repeating square-wave glucose perturbations, studying the effects on growth and composition of *E.coli* ML30. However, they applied cycles of 2 hours duration, which are not able to capture the metabolic responses occuring in timescales of seconds in large-scale bioreactors. Sunya S*, et al.* [39] characterized the dynamic behaviour of *E. coli* DPD2085 imposed in 4 successive cycles, of 7 min duration each, of glucose pulses in different intensities. All cycles were compared to each other, in terms of specific formation and consumption rates. Nonetheless, 4 cycles were not enough to obtain repetitive O_2_ and CO_2_ concentration profiles, indicating that the microorganism was not yet physiologically adapted. In addition, the intracellular metabolic activity was not monitored.

In this study, a feast-famine regime was applied, for the first time, in a physiologically adapted *E.coli* K12 aerobic culture, with successive glucose perturbations and both intracellular and extracellular metabolic responses, occuring in short-time scale (seconds), were quantitatively described.

## 2. Materials and Methods

### 2.1 Strain and cultivation medium

*Escherichia coli* wild-type strain K12 MG1655, obtained from The Netherlands Culture Collection of Bacteria (NCCB, 3508 AD, Utrecht, The Netherlands), was used in all the experimental work of this study. The cultivation (and preculture) medium consisted of (per litre): 0.151 mol glucose (C_6_H_12_O_6_·1H_2_O), 0.5 g MgSO_4_·7H_2_O, 0.5 g NaCl (Avantor J.T.Baker, Gliwice, Poland), 1.25 g (NH_4_)_2_SO_4_, 1.15 g KH_2_PO_4_, 6.75 g NH_4_Cl (Merck KGaA, Darmstadt, Germany), 0.001 g thiamine-HCl (Sigma-Aldrich, St. Louis, Missouri, USA) and 2 mL of trace elements solution [40]. The pH of the medium was adjusted to 7.0 by the addition of 1 M K_2_HPO_4_, before filter sterilization (pore size 0.2 μm, cellulose acetate, FP 30/0.2, Whatman GmbH, Dassel, Germany).

### 2.2 Preculture

Culture aliquots, previously stored in 80% v/v glycerol at −80 °C, were used for the preculture. Cells were grown in shake-flasks filled with 100 mL of the above-mentioned mineral medium, in an incubator (37 °C, 220 rpm) and were used as inoculum for the bioreactor cultivation.

### 2.3 Cultivation conditions

The cultivation was performed in a 1.2 L stirred tank bioreactor (Applikon Biotechnology B.V., Delft, The Netherlands), with 0.95 L working volume, controlled by weight. The bioreactor was aerated with pressurized air at 0.44 L min^-1^ (0.5 vvm), using a Smart series mass flow controller 5850S (Brooks Instrument, PA, USA). The bioreactor was operated at 0.3 bar overpressure, at 37 °C and a stirrer speed of 700 rpm. pH was controlled at 7.0 by automatic addition of either 4 M KOH or 2 M H_2_SO_4_. Antifoam (Basildon Chemicals Ltd, UK) was added manually, when necessary, during the batch phase. During the whole experiment, pH, temperature, medium and effluent vessel weight, base and acid addition were monitored online. In addition, the dissolved oxygen in the broth was measured by a polarographic ADI sensor (Applisens, Applikon, Delft, The Netherlands). A gas analyser (NGA 2000, Rosemount, Emerson, USA) was used to measure on-line the oxygen (O_2_) and carbon dioxide (CO_2_) concentrations in the offgas. After the batch phase was completed (indicated by the decrease of carbon dioxide and the increase of dissolved oxygen), the medium feeding was switched on and a chemostat phase began at a dilution rate of 0.044 h^-1^. After 114 hours (5 residence times), samples for metabolite and biomass quantification were withdrawn.

### 2.4 Dynamic feast-famine regime

After sampling of the reference chemostat, the regime was changed to an intermittent feeding. The feast-famine setup is shown in Figure 1. Successive cycles of 400 s were applied by a continuous medium feeding for 20 s, followed by a period of 380 s of no feeding. The feeding pump was controlled automatically by a timer (Omega, CT, USA). The waste outflow was controlled by a scale, on top of which the reactor operated, maintaining the broth volume at 0.95 L. The regime was designed to feed the same amount of medium over time, as in the chemostat culture, leading to an average dilution rate of 0.047 h^-1^. Aeration rate was increased to 0.7 L min^-1^ (0.8 vvm), to avoid oxygen limitation. The rest of the cultivation conditions remained the same as the reference chemostat. After 180 hours (8 residence times) of the intermittent feeding, samples for biomass and metabolite quantification were withdrawn. Sample analysis

**Figure 1.**
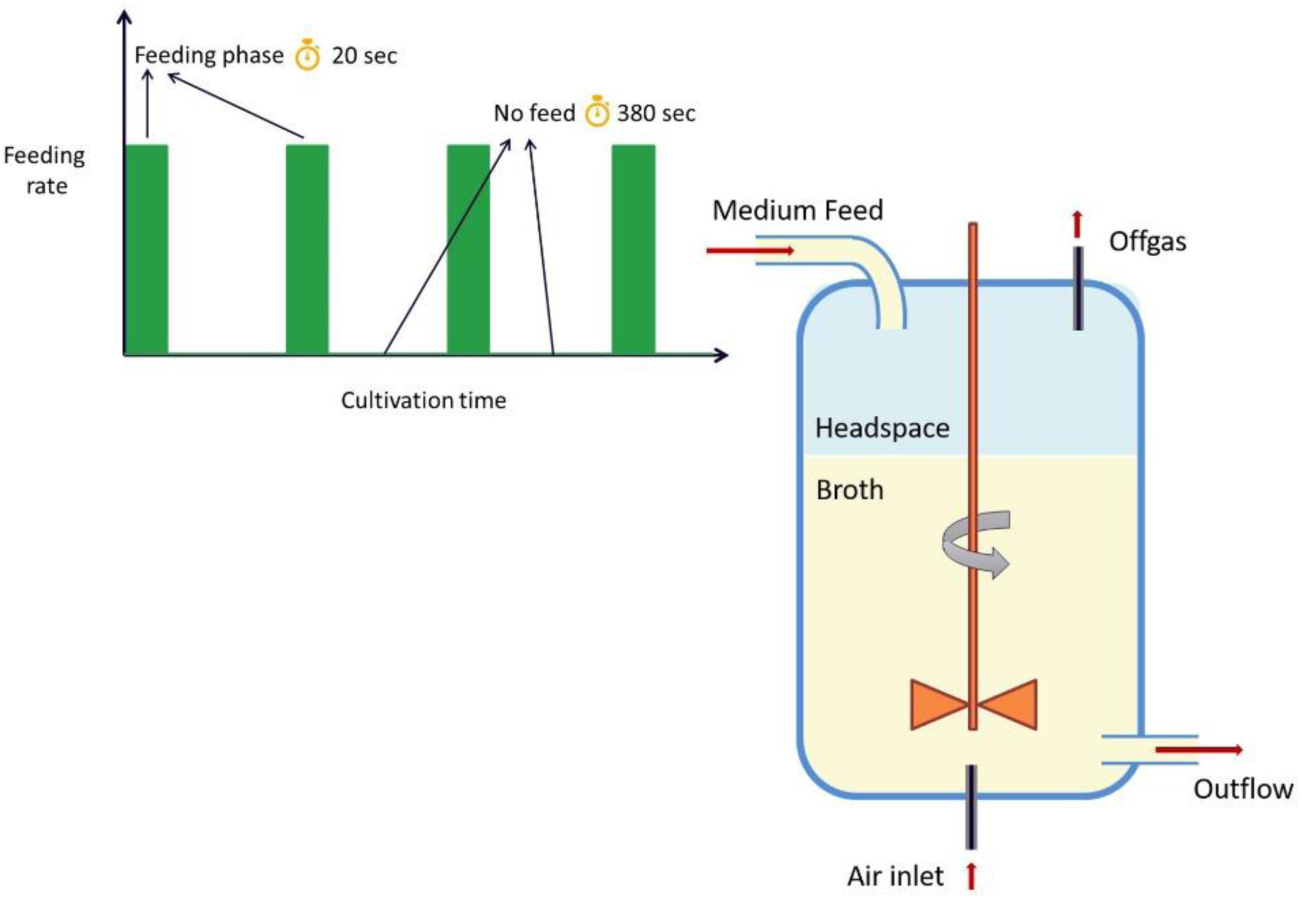
Schematic representation of the feast-famine setup used in this work. The medium, containing glucose as a substrate, was fed block-wise (20 s on, 380 s off) through a head-plate port. A constant volume was maintained by weight control. Successive cycles run for about 200 h in total.

#### 2.4.1 Cell dry weight measurements

For the determination of the biomass concentration (dry weight), 2 mL of broth were collected and centrifuged (Heraeus Biofuge Stratos centrifuge) at 4 °C for 5 min at 13800 g. The supernatant was then discarded and the pellet was resuspended in 1 mL Milli-Q water. Centrifugation and resuspension were repeated, the sample was then transferred to a previously dried (for 48 h at 70 °C) and weighted glass vial and dried at 70 °C for at least 48 h. The vials were then weighted again (after cooling down to room temperature inside a desiccator) and the cell dry weight was calculated as the difference between the final weight and the empty vial weight. The average of four replicate samples was used for the steady-state culture, six replica were used for the feast-famine regime.

#### 2.4.2 Total organic carbon

Total broth samples (2 mL each) were withdrawn from the reactor and immediately stored at −80 °C. Supernatant samples were acquired by centrifuging total broth samples (Heraeus Biofuge Pico microcentrifuge) at room temperature for 1 min at 10400 g. The supernatant was then filtrated (pore size 0.2 μm filter, cellulose acetate, FP 30/0.2, Whatman GmbH, Dassel, Germany) and stored at −80 °C. The total amount of organic carbon (TOC) was quantified with a TOC Analyzer (TOC-L CSH, Shimadzu), using the “difference method”: TOC was calculated from the difference between total carbon and inorganic carbon. Calibration standards were obtained from LPS b.v. (Oss, The Netherlands).

#### 2.4.3 Extracellular metabolite sampling

For the determination of extracellular metabolite concentrations, 2 mL of broth were withdrawn into a tube (eppendorf) and immediately centrifuged (Heraeus Biofuge Pico microcentrifuge) at room temperature for 1 min at 10400 g. The supernatant was then filtrated (pore size 0.2 μm filter, cellulose acetate, FP 30/0.2, Whatman GmbH, Dassel, Germany) into an empty tube, submerged into liquid nitrogen and stored at −80 °C until analysis. Centrifugation was used before filtration to prevent blocking of the filter, due to the high biomass concentration of the samples. For GC-MS and LC-MS analysis, 60 μL of ^13^C cell extract were added to 300 μL of sample, as internal standard mix, before freezing with liquid nitrogen and storage at −80 °C until further analysis.

#### 2.4.4 Intracellular metabolite sampling

For the determination of intracellular metabolite concentrations, the differential method was applied with some modifications [41]. For the total broth measurement, 1 mL of broth was withdrawn from the reactor into a tube filled with 5 mL aqueous methanol quenching solution (60% v/v) at −40 °C, to rapidly stop metabolic activity. The sample was immediately vortexed to ensure homogeneity and then weighted. 120 μL of ^13^C cell extract (production method described in [42]) were added to the sample, as internal standard mix. For the extraction of metabolites, 5 mL of aqueous ethanol solution (75% v/v), preheated at 70 °C, were added to the sample and the tube was then placed into a water bath at 95 °C for 4 minutes. After the boiling extraction, the sample was immediately cooled down to −40 °C in a cryostat.

The ethanol-water mixture in all samples was then evaporated in a Rapid-Vap (Labconco, MO, USA) at 30 °C, under vacuum. The dried sediment was resuspended in 600 μL Milli-Q water, vortexed and transferred to eppendorf tubes. The samples were centrifuged at 15000 g for 5 minutes at 1 °C (Heraeus Biofuge Stratos centrifuge). The supernatants were transferred to new empty tubes and centrifuged again under the same conditions. The filtrate was stored in screw-cap vials, at −80 °C, until further analysis. The intracellular concentrations were obtained from the difference between total broth and extracellular measurements.

#### 2.4.5 Analytical methods

Extracellular concentrations of organic acids (acetate, lactate, formate) and ethanol (from 2.4.3) were determined by HPLC (BioRad HPX-87H 300*7.8 mm column, at 59 °C, 0.6 mL·min^-1^, 1.5 mM phosphoric acid in Milli-Q water as eluent, coupled to a Waters 2414 RI detector and a Waters 2489 UV detector at 210 nm).

Processed extracellular and total broth samples were analysed by GC-MS/MS, GC-MS and LC-MSMS. Metabolites of the central carbon pathways (glycolysis, pentose phosphate pathway (PPP), tricarboxylic cycle (TCA)) were quantified with GC-MS/MS (7890A GC coupled to a 7000 Quadrupole MS/MS, both from Agilent, Santa Clara, CA, equipped with a CTC Combi PAL autosampler, CTC Analytics AG, Zwingen, Switzerland), as described in [43] and/or anion-exchange LC-MSMS [44]. GC-MS was used for the quantification of amino acids, as described in [45]. Ion-pair reversed phase LC-MSMS was used for the quantification of nucleotides, as described in [46]. The isotope dilution mass spectrometry (IDMS) method, described in [42, 47], was used for the metabolite quantification.

## 3. Results

The adaptation of *Escherichia coli* to repetitive, dynamic perturbations was evaluated with respect to short and long-term physiological and metabolic characteristics:

1. Comparison between average metabolic rates and yields under repetitive dynamic conditions and steady-state (reference) conditions, at the same dilution rate.
2. Comparison of the metabolic response between repetitive dynamics, single perturbations (pulse experiments) and steady-state levels.

It was assumed that cells were adapted after five residence times under the feast-famine regime, and a repetitive metabolic response was obtained. This assumption was supported by the observation that the online measurements of dissolved oxygen (DO) and offgas (O_2_ and CO_2_) concentrations showed a highly reproducible pattern over the cycle (Figure 2).

**Figure 2.**
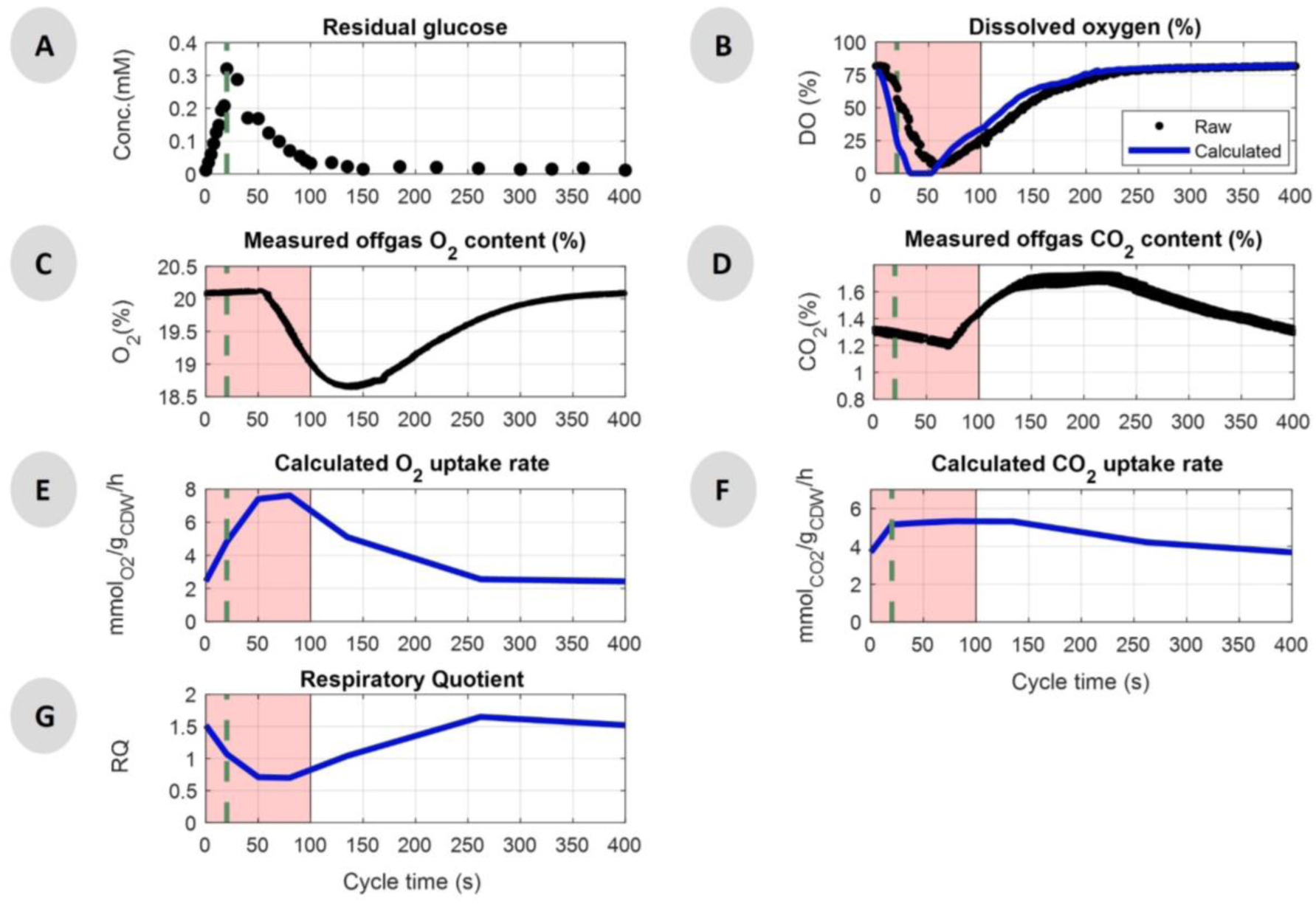
Measured concentrations and calculated rates during the feast-famine cycle (s), approximately 8 generations after the beginning of the regime. A) Residual glucose concentration (mM), quantified by GC-MS/MS. B) Dissolved oxygen concentration (%) in the broth, raw data (black) and calculated, eliminating delays of the used Clark probe, (blue). C) Measurements of oxygen content in offgas (%). D) Measurements of carbon dioxide content in offgas (%). E) Calculated oxygen uptake rate (mmol_O2_/g_CDW_/h) based on the headspace and tubing offgas delays. F) Calculated carbon dioxide production rate (mmol_CO2_/g_CDW_/h) based on the headspace, tubing offgas delays and bicarbonate in the broth. G) Respiratory quotient (RQ) over time derived from the calculated q_O2_ and q_CO2_. Data of 16 successive cycles are overlapped for DO, O_2_ and CO_2_ (B, C, D). The pink area in the plots represents the substrate feast phase. Green vertical dashed lines show the end of the feeding (20 s).

### 3.1 *Escherichia coli* physiological behaviour under substrate dynamics

#### Extracellular environment

During the first 100 seconds of the cycle the residual glucose concentration (Figure 2A) was higher than the reported glucose affinity constant (*K_M_* = 10 μM) of the microorganism [48]. This time period will be referred to as (substrate) feast phase. The concentration increased from 0.01 to 0.32 mM, which was the maximum value, during the first 20 s of the cycle. Then, glucose depleted after 100 s (< 10 μM), leading to a (substrate) famine phase.

The broth dissolved oxygen profile was estimated by deconvolution (Figure 2B blue line), i.e. accounting for the dynamics of the Clark electrode (see Supplementary Material S1 for details). With the supply of substrate, a decreased dissolved oxygen concentration was observed, suggesting a (high) oxygen consumption during the substrate feast period. The same behaviour was expected for the offgas measurements. The minimum O_2_ concentration (and maximum CO_2_) was observed only after the end of the feast phase. This delay can be explained by the headspace and tubing gas hold-up [49]. The O_2_ uptake rate and CO_2_ production rate over time (Figure 2 (E-F)) were therefore calculated, taking into account these delays (Supplementary Material S2). For the CO_2_ production, the interconversion of dissolved carbon dioxide to bicarbonate in the broth, due to the neutral pH 7 was, also, taken into account. The respiratory quotient was then derived from these rates over time (Figure 2G). We observed that the RQ decreased from 1.5 to 0.7 during the first 50 s of the feast phase, and increased back to the initial value after approximately 200 s, indicating that the electrons, from 20 to 120 s of the regime (RQ < 1), are transferred to oxygen, while the respective carbon is not found in the form of CO_2_. Therefore, we expect a form of intracellular storage compound (with electron to carbon ratio e^-^/C < 3), synthesized in the feast phase and degraded in the famine phase. Further discussion on this follows after the metabolome section (3.3).

#### Average biomass specific rates and yields

In order to compare the physiology of the cells between steady-state and feast-famine conditions, the respective biomass specific conversion rates were calculated (Table 1). For the steady-state culture, rates were derived from the respective mass balances. During the intermittent feeding regime, all of the glucose supplied over one cycle was consumed, according to the GC-MS/MS measurements. Therefore the average glucose uptake rate (q_glc_) was calculated as the amount of substrate fed over the total cycle time. Taking into consideration the 20 h average doubling time of *Escherichia coli* at a dilution rate of 0.05 h^-1^, we expected the biomass concentration to vary by 0.5% during one cycle. This small change could not be determined experimentally, therefore a constant specific growth rate (μ) was considered within a cycle and calculated using the average of six cell dry weight measurements (see 2.4.1). To calculate the average O_2_ uptake and CO_2_ production rates, the respective online measurements were first integrated (trapezoidal method) over time, and then averaged for 16 successive cycles.

**Table 1.** Steady-state and average feast-famine biomass specific rates with their associated standard deviations. All the results presented in the table were calculated using data reconciliation. Raw data can be found in Supplementary Material S4.

|  | Steady-state | Feast-famine<br>(cycle average) | log2-fold<br>changes between |
| --- | --- | --- | --- |
| **Viable Biomass Concentration (g L<sup>-1</sup>)** | 9.33 ± 0.01 | 6.28 ± 0.03 | - 0.6 |
| **Biomass Growth $\mu$ (g g<sub>CDW</sub><sup>-1</sup> h<sup>-1</sup>)** | 0.044 ± 0.002 | 0.048 ± 0.003 | + 0.1 |
| **Lysis rate (g g<sub>CDW</sub><sup>-1</sup> h<sup>-1</sup>)** | - | 0.008 ± 0.006 | - |
| **$q_{\text{Glucose}}$ (mmol<sub>glc</sub> g<sub>CDW</sub><sup>-1</sup> h<sup>-1</sup>)** | -0.73 ± 0.01 | -1.12 ± 0.02 | + 0.6 |
| **$q_{\text{O}_2}$ (mmol<sub>O2</sub> g<sub>CDW</sub><sup>-1</sup> h<sup>-1</sup>)** | -2.08 ± 0.02 | -4.42 ± 0.06 | + 1.1 |
| **$q_{\text{CO}_2}$ (mmol<sub>CO2</sub> g<sub>CDW</sub><sup>-1</sup> h<sup>-1</sup>)** | 2.30 ± 0.02 | 4.53 ± 0.06 | + 1.0 |
| **Respiratory Quotient** | 1.11 ± 0.01 | 1.03 ± 0.02 | - 0.1 |
| **Total Biomass Yield (g<sub>CDW</sub> g<sub>glc</sub><sup>-1</sup>)** | 0.31 ± 0.01 | 0.18 ± 0.03 | - 0.5 |
| **Oxygen to Substrate ratio** | 2.85 ± 0.05 | 3.94 ± 0.08 | + 0.5 |

The extracellular by-product (acetate, lactate, formate and ethanol) concentrations (Supplementary Material S3), obtained from HPLC measurements, were also integrated over the cycle and their average formation rates were then calculated from their mass balances (raw data can be found in Supplementary Material S4). The average acetate production did not exceed 0.02 mmol/g_CDW_/h, both during steady-state and feast-famine conditions. The absence of high amounts of by-products in *E.coli* cultivations has been observed in previous studies, both during chemostat (usually at lower dilution rates (up to 0.3 h^-1^)) and after glucose or oxygen perturbations [50–54]. The by-product formation is related to overflow metabolism, a common phenomenon in *E.coli* cultivations in glucose-containing medium, where acetate is produced even when the culture is fully aerated [55–57]. Here, overflow metabolism was not significantly affected by the transition of the cells from steady-state to long-term dynamics.

The biomass specific rates were reconciled using the approach described in [58], using element conservations as constraints and the total organic carbon (TOC) measurements of the broth and the filtrate. A constant biomass composition of CH_1.73_N_0.24_O_0.35_S_0.006_P_0.005_ (γ_Χ_=4.38, M_W_=23.2 g/Cmol) [59] was used for all calculations. The phenomenon of cell lysis was also introduced in the reconciliation to account for carbon and electrons of non-viable biomass.

The most evident observation is that the calculated rates for glucose and oxygen uptake and CO_2_ production were higher (+0.6, +1.1, +1.0 log2-fold times respectively) for the intermittent feeding, compared to the reference continuous regime. More specifically, during the reference chemostat operation, the oxygen to substrate consumption ratio was 2.85 ± 0.05 mol_O2_/mol_glc_, which is comparable to the values reported in literature for similar cultivation conditions, ranging from 2.74 – 3.62 mol_O2_/mol_glc_ [50, 51, 59–61]. When the cells were subjected to feast-famine cycles, the average oxygen to substrate ratio increased, reaching 3.94 ± 0.08 mol_O2_/mol_glc_.

Introducing potential cell lysis in the reconciliation, we observed that the lysis rate during the steady-state was calculated as negative (while positive by definition), with a high standard deviation and therefore the steady-state lysis was assumed zero.

The intermittent feeding, resulted in decreased average cell dry weight concentration (viable biomass) and subsequent 0.5 log2-fold decrease (−30.3 %) of the total biomass yield. The biomass yield was calculated as the ratio of the total biomass growth rate (including lysed biomass) over the glucose uptake rate. A decrease in biomass yield has also been reported in previous studies for *E.coli,* when varying the substrate availability. Many studies attributed this phenomenon either to overflow metabolism, due to high growth rates (>0.15 h^-1^) [25, 39], or to oxygen limitation. For example, Neubauer P*, et al.* [62] observed a 10% decrease in yield for cells circulating from zones of glucose excess to glucose starvation in a scale-down two-compartment reactor (continuous stirred and plug flow), compared to reference conditions. This was explained by the oxygen limitation occurring in high-concentrated regions of glucose. However, this was not the case in our experimental setup. Only minor amounts of acetate were produced as a by-product and did not increase under the dynamic conditions. In addition, we did not observe any changes in the biomass composition during the reference conditions compared to the one during the feast-famine regime (elemental analysis, Supplementary Material S5). Lower biomass yields were also reported for other microorganisms, such as *Saccharomyces cerevisiae* (almost 25% decrease), subjected to an intermittent feeding (similar to the regime used here) [63].

The ratio of CO_2_ produced per glucose consumed, during the steady-state regime, was 3.16 ± 0.05 mol_CO2_/mol_glc_, while for the feast-famine a value of 4.05 ± 0.08 mol_CO2_/mol_glc_ (+ 28.1%) was observed. Together with the increase in O_2_ consumption, it indicates that more glucose was used for respiration, rather than biomass production, during the feast-famine cycles [8].

### 3.2 Glucose uptake dynamics: a matter of seconds

The feast-famine setup allows to measure the short-term responses of *E.coli* cells with high time resolution and repeated measurements. Especially, samples can be obtained from repetitive cycles (Figure 2B, C, D), which allowed for distributing sampling over several cycles.

Special focus was to obtain a high-resolution uptake profile as substrate uptake has a major impact on the intracellular metabolic behaviour of the microorganism. To calculate the short-term glucose uptake profile, a piecewise affine (PWA) rate approximation [64] was calculated. The breakpoints used were timepoints of 0, 2, 15, 18, 110 and 400 s. These breakpoints were chosen based on the highest goodness of fit (R^2^ was used), among various combinations [65]. The flux between the breakpoints followed a first order linear function (Figure 3). Note that the first and last breakpoint were coupled (cyclic regime). The rate was normalized using the reconciled biomass concentration of 6.28 g L^-1^.

**Figure 3.**
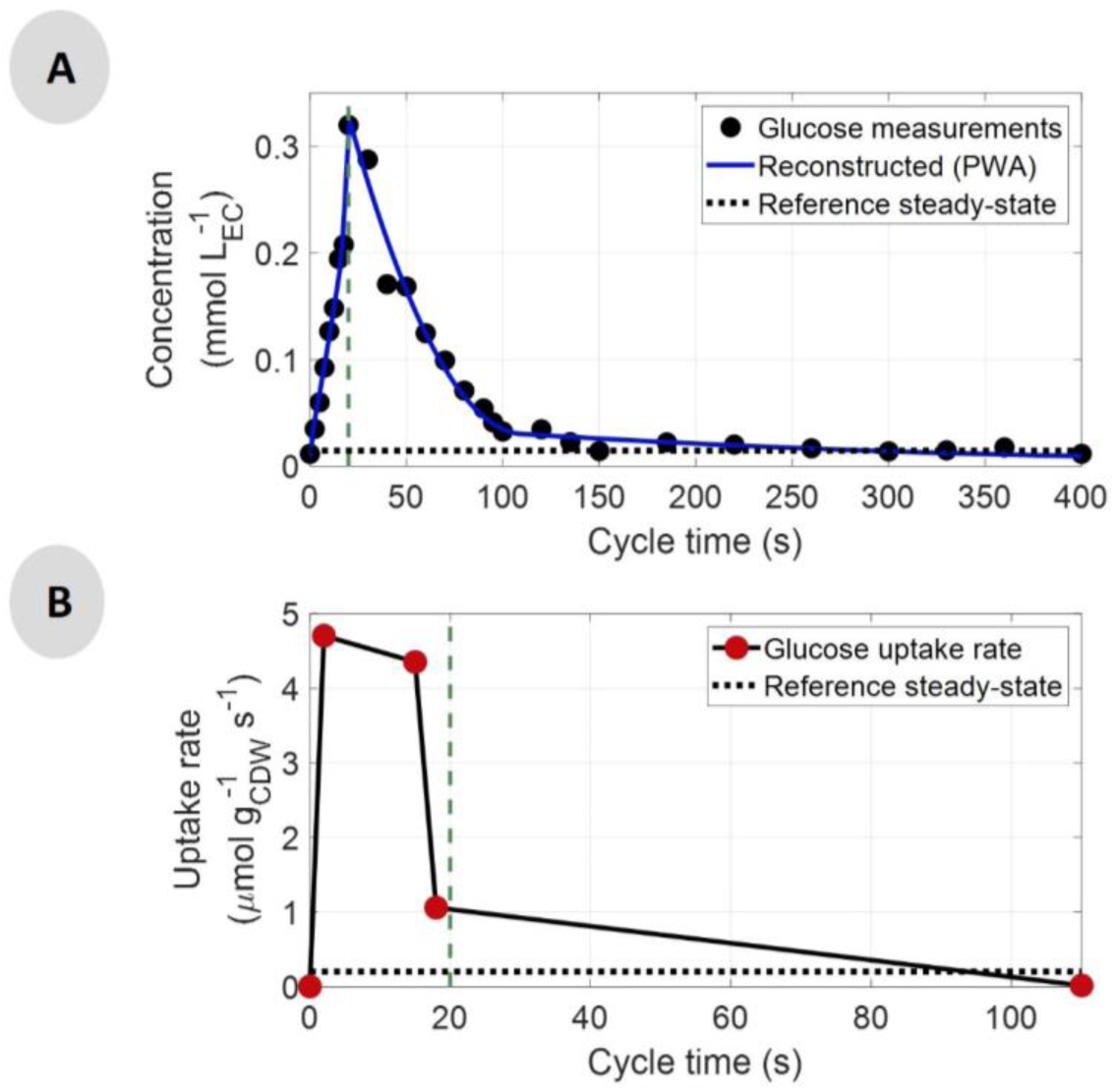
A) Residual glucose concentration in mM (black dots) and the PWA fitted profile (blue line) over one cycle time (s). B) Glucose uptake rate (-q_glc_) in μmoles_glc_/g_CDW_/s over one cycle time (s). The red dots represent the breakpoints. Note that for the rate only the first 110 s are shown. For t > 110s, the flux was zero until the end of the cycle. Green vertical dashed lines show the end of the feeding (20 s) and the horizontal dotted lines represent the average steady-state levels.

There was no obvious correlation between the glucose uptake rate and the extracellular glucose concentration. The rate reached its highest value (4.68 μmol/g_CDW_/s) immediately after the beginning of the feeding and then decreased slightly until the end of the feeding, followed by significant decrease between 15 s and 18 s, i.e. from 4.40 to 1.04 μmol/g_CDW_/s.

This contradiction is explained by the activity of the phosphotransferase system (PTS) [66], which is the main substrate uptake system in *E.coli* under glucose excess (Km is in the range of 3-10 μΜ [48, 67]). Therefore, the glucose transport did not depend only on the extracellular concentration, but also on the intracellular concentrations of other metabolites like G6P, PEP and pyruvate, which are key components of the PTS.

The overshoot in the glucose uptake rate has been previously observed in cells exposed to excess of substrate after a starvation period, in different experimental setups [52, 60, 62]. However, the main difference of our work is that we described an adapted microorganism, which has sustained substrate perturbations for more than 8 generations, while the above-mentioned studies reported the behaviour of the cells right after applying a perturbation for the first time.

The highest estimated uptake rate in our work was significantly higher than the maximum observed during batch cultivations for the same strain (3.06 [68] and 3.30 [52] μmol/g_CDW_/s), proving that the microorganism has a higher uptake capacity than the one observed under maximum growth. It is, however, puzzling that the glucose uptake rate was decreasing already (mainly after 18 s), while glucose was still in excess (above the glucose affinity constant). An additional test experiment was performed under the same cultivation conditions, with a shorter feeding phase (13 s) but the same cycle length (400 s). The glucose uptake rate calculated (data not shown) exhibited the same decreasing pattern 2 seconds before the end of the feeding phase (at 11 s), while glucose was still in excess. Therefore, it is concluded that the specific feeding time, chosen for the experimental setup of this study, was not an influencing factor of this behaviour. This decrease suggests that there was another limitation in further metabolizing glucose, which arose early in the feast phase. Another critical observation is that the uptake of glucose was decoupled from the oxygen uptake. While there was no glucose uptake after 100 s (Figure 3B), oxygen was still consumed (Figure 2Ε), suggesting that an intracellular compound was oxidised during the substrate famine phase. Different hypotheses for both observations will be derived (see Discussion) based on the analysis of the intracellular metabolite measurements (section (3.3)).

### 3.3 Metabolite dynamics in the intracellular space

The highly dynamic glucose uptake rate, discussed in section 3.2, was expected to result in significant fluctuations in the intracellular metabolite levels. Therefore, it is important to observe how *E.coli* is regulating its metabolic network to handle the increased fluxes under these rapidly changing conditions, without detrimental effects on its survival. For this aim, a broad range of intracellular metabolites were quantified during the reference chemostat regime and the feast-famine cycle regime, including glycolysis, tricarboxylic acid (TCA) cycle, pentose phosphate pathway (PPP) intermediates and amino acids.

#### Glycolysis and Pentose Phosphate Pathway

The intracellular flux profiles over time were estimated by (dynamic) Flux Balance Analysis (FBA) for the glycolytic and pentose phosphate reaction steps. The stoichiometric network included reactions of central carbon metabolism, where the intermediates could be measured (otherwise lumped, see Supplementary Material S6 for details). The glucose uptake rate from section 3.2 was used as input for the PTS flux. For the steady-state flux determination it was assumed that 70% of the PTS flux was directed towards glycolysis, while the rest was directed towards the pentose phosphate pathway; this ratio is a recurring value reported in literature [59, 69–71]. For the dynamic flux determination during the feast-famine cycles, instead of setting the PTS flux split ratio, we used the minimization of the squared difference between the dynamic and the steady-state PGI and PPP fluxes as the optimization target (i.e. minimization of metabolic adjustment – MOMA) [72]. It has been suggested that mutant *E.coli* strains redistribute their metabolic fluxes in such way to minimally divert from the wild-type metabolic network [72–74]. Our work is a comparable case, where the perturbation of the cells was not genetic, but kinetic, as a result of an intermittent substrate feeding. It was, therefore, assumed for our analysis that the optimal flux distribution after the perturbation required the smallest change from the steady-state metabolism, the same way the genetic engineered strains adapt with respect to the wild-type.

The simple network used in FBA is shown in Figure 4, along with the measured metabolite concentrations and the estimated fluxes over time. MOMA was performed in Matlab R2018a, The MathWorks, Inc., using quadratic programming. The derived fluxes in the chosen breakpoints can be found in Supplementary Material S7.

**Figure 4.**
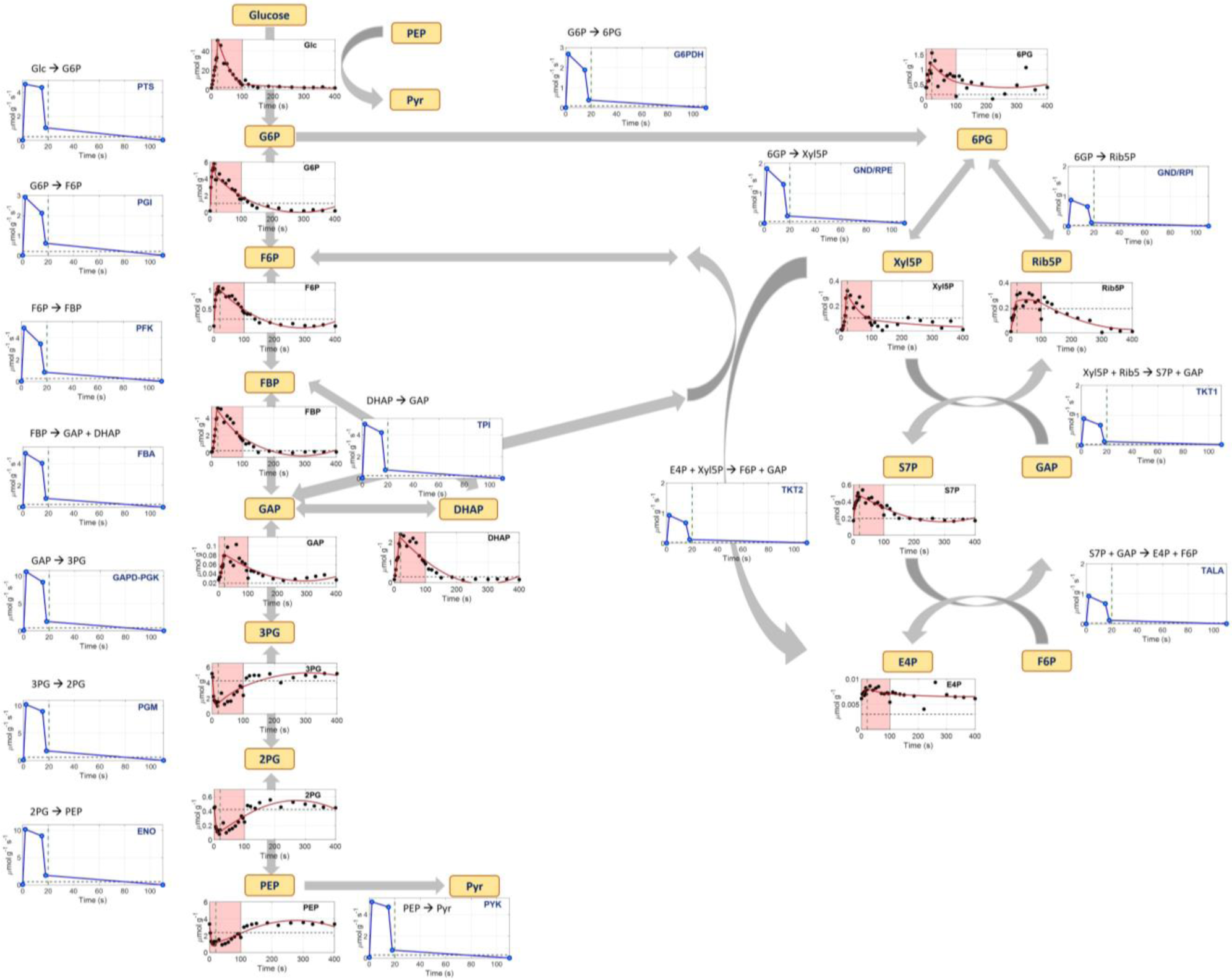
Network model used for the flux balance analysis. Metabolite names are shown in yellow text boxes. Under each metabolite, its intracellular concentration (μmol/g_CDW_) (extracellular only for glucose) over time (s) is shown. Black dots represent the measurements, the red line is the PWA fitted line and black dashed lines represent the average steady-state levels. Green vertical dashed lines show the end of the feeding (20 s). The pink area represents the substrate feast phase. The blue line plots show the FBA estimated flux profiles in μmol_reaction_substrate_/g_CDW_/s, where the blue dots are the values at the breakpoints. Fluxes are shown up to 110 seconds and they were all zero afterwards until the end of the cycle.

Looking at the dynamics in terms of concentration profiles, we observed that the microorganism transported the glucose from the extracellular to the intracellular space, causing all the upper glycolytic metabolites to increase during the first 20 s, in agreement with the extracellular glucose decrease. After fast filling of the pools, depletion also followed the extracellular concentration profile, i.e. very low concentration levels were observed during the famine phase (> 100 s). The measured concentrations were significantly higher than the steady-state levels (e.g. up to 20 fold change for FBP), due to the short-term overshoot of carbon flux. The opposite trend was observed for the lower glycolytic metabolites (3PG, 2PG, PEP), whose concentration immediately decreased in the first seconds of the feast phase (3-6 fold change from steady-state). The reason behind this drop is based on the PTS system. In order to import and phosphorylate glucose to G6P, PEP needs to be produced and then converted to pyruvate. Unfortunately, pyruvate could not be quantified in this experiment. PEP concentration showed negative correlation with G6P, as it reached its lower concentration after 12.5 s, the same time the maximum concentration of G6P was reached. This behaviour has been observed before in *E.coli* responses to glucose pulses [50, 75–77]. Even in lower concentrations PEP was always available, during the whole cycle, for the import of glucose, therefore no limitation in the glucose transport system of *E.coli* was observed.

The metabolites of pentose phosphate pathway related well to the dynamics observed in glycolysis, as they exhibited the same behaviour of rapid accumulation in the beginning of the feast phase and later decrease to the initial levels. The pool of 6PG responded directly to the changes occurring in its precursor, G6P, reaching its maximum concentration at 20 s (7.5 s delay compared to G6P). This peak was observed slightly later in the rest of the metabolites, with the exception of Xyl5P, which responded equally fast.

Looking at the flux profiles, we observed that the glucose uptake rate dynamics propagated through glycolysis. The peak observed in the PTS flux right at the beginning of the feeding, also occurred in the succeeding reactions towards the formation of PEP and they all decreased significantly after 15 s. After 110 s all metabolite pools remained constant, while there was no more flux running in glycolysis. The same trend was also observed in all the reaction steps of the pentose phosphate pathway.

Compared to the steady-state levels, the immediate increase of the PTS flux (16 fold) led to a higher change in all the glycolytic fluxes (18-19 fold) and an even higher increase in the flux towards PPP (30 fold). This observation, together with the fact that 62.4% of the PTS flux was directed into glycolysis (less than the 70% assumed in steady-state), gives an indication that the cells may increase the flux to pentose phosphate pathway, under these dynamic conditions, therefore enhancing the production of NADPH, assumingly for redox balance purposes. Similar increase was observed after a single-pulse of glucose in an aerobic *E.coli* culture, which was used to further support the calculated increase in growth rate during the feast phase [52]. NADPH was, therefore, needed to support the increased growth. This behaviour has, also, been observed as a response to oxidative stress for *E.coli* [78, 79] and other organisms [80, 81].

#### TCA Cycle

In the case of the TCA cycle, only a few metabolites could be precisely quantified, which are shown in Figure 5.

**Figure 5.**
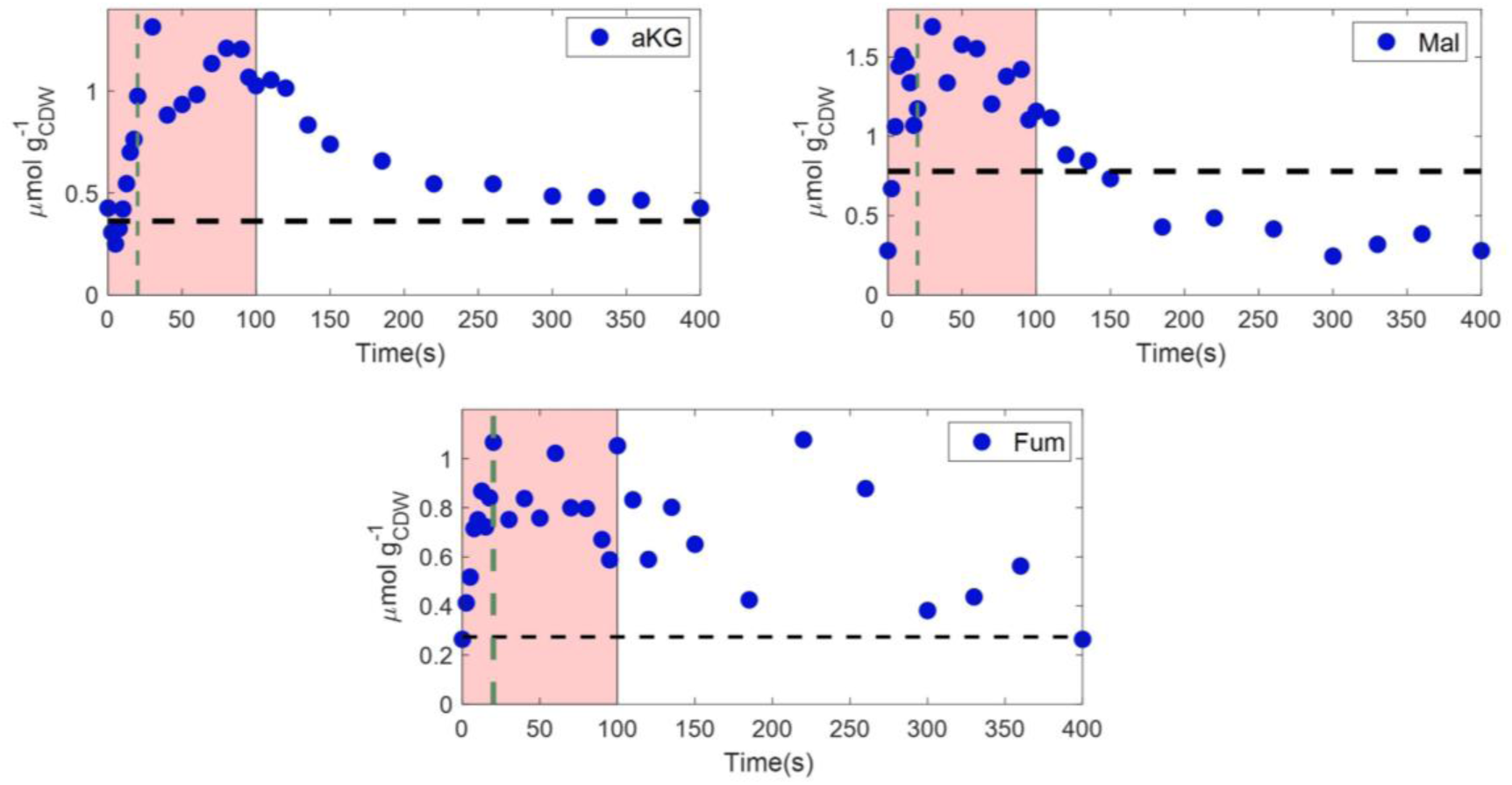
Intracellular concentrations (μmol/g_CDW_) of TCA metabolites (a-ketoglutarate, malate and fumarate), over a feast-famine cycle (s). Black horizontal dashed lines represent the average steady-state levels. Green vertical dashed lines show the end of the feeding (20 s).The pink area represents the substrate feast phase.

Following glycolysis, also the TCA metabolites showed a dynamic profile over time. We observed that aKG and malate reached their highest concentration 30 s after the beginning of the feast-famine cycle (Figure 5), displaying a delay of 10 s compared to the glucose profile (Figure 3). This delay was also evident during the famine phase, as the metabolite levels reached low levels much later than the glycolytic ones (>100 s). Fumarate showed a more oscillating profile over time, but a general increase of the pool during the feast phase and a following decrease, to the initial levels at the end of the cycle, was detected. The highest concentration change during the cycle was observed for malate (6.1 fold), which was still lower than the dynamics of the upper glycolytic metabolites, such as G6P (31.7 fold) and F6P (18.8 fold).

#### Amino acids

Amino acids are relevant precursors for protein synthesis. At the same time, several amino acids are closely connected with their respective central carbon metabolism precursor. For example alanine is only one (equilibrium) reaction step from pyruvate. Thus, on the one hand one could expect homeostasis to ensure balanced growth, on the other hand a high dependency on central carbon metabolism (Supplementary Material S8). Amino acids derived from E4P (Supplementary Material S8 – Figure S.1) displayed a similar dynamic profile with their precursor, with delays in reaching their highest values. The same trend was observed for the amino acids derived from aKG (Supplementary Material S8 – Figure S.5) and the ones from pyruvate. They all increased and decreased over time related to the concentration of their precursors, with some exhibiting more pronounced and faster dynamics (e.g. glutamine, alanine, leucine) than others (e.g. lysine, proline). On the other hand, serine, tryptophan and glycine were not significantly affected by the profile of their precursor, 3PG, as they displayed small changes over time, remaining close to their steady-state values (Supplementary Material S8 – Figure S.2). The amino acids, derived from oxaloacetate, also, displayed various trends, either rapidly increasing (e.g. threonine) or decreasing (e.g. aspartate, cysteine) during the feast phase (Supplementary Material S8 – Figure S.3).

The largest deviation of concentrations from the steady-state ranged from 1 to 3 fold times for most of the amino acids, more modest than the changes in their precursors. Cysteine was the only exception, as its concentration was measured to be 200 fold higher than the steady-state, in the beginning of the feast-famine regime (Supplementary Material S8 – Figure S.3). Interestingly, while all amino acid concentrations decreased at the end of the cycle towards biomass synthesis, some even reaching their low steady-state levels, the opposite trend was observed for aspartate. Aspartate decreased rapidly during the feast phase and then increased during the famine phase, reaching a concentration of around 6 μmol/g_CDW_, while its steady-state concentration was 2.6 μmol/g_CDW_ (Supplementary Material S8 – Figure S.3).

#### Nucleotides and Energy Homeostasis

Nucleotide responses to the feast-famine regime are of high interest and especially the ATP/ADP levels, which reflect the adenylate energy state (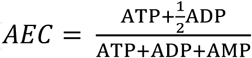 [82]) of a cell and can provide insights on how the cells encounter the dynamic perturbations energetically. As expected, but still surprising, the AEC of the cell showed stability throughout the cycle (average value of 0.79), indicating that the total rate of ATP production is equal to the one of ATP consumption. The average AxP (sum of ATP and ADP) concentration was in the range of 6.99 μmol/g_CDW_, while the energy turnover is expected in the range of 0.4 to 1 second, using a normal P/O ratio (2.98) for *E.coli* [59]. Therefore, such balancing occurred in sub-seconds. Under glucose-limited and batch growth conditions the AEC, in most microorganisms, ranges between 0.7-0.95 [83–86]. AMP concentration was at noise level and therefore not quantified. Its contribution on the AEC calculations was neglected.

Our findings are in agreement with results from single pulse experiments [50, 52, 87], as well as two-compartment scale-down cultivations [34], in *E.coli* K12, where the energy homeostasis was also reported during glucose excess (AEC ranging from 0.8 to 0.85 in the different studies). However, in these studies, the AEC was decreasing during the famine phase, sometimes reaching values even lower than 0.7 [34], which was not the case in our experiment. Link H*, et al.* [88] also observed that the AEC remained unaffected, ranging between 0.7-0.8, after transferring fed-batch grown cells to batch reactors.

In contrast to AxP’s, UDP and UTP were not homeostatic. They decreased over the feast phase and increased during the famine phase. Also, GDP and GTP were dynamic with a trend opposite to UxP’s (Figure 6).

**Figure 6.**
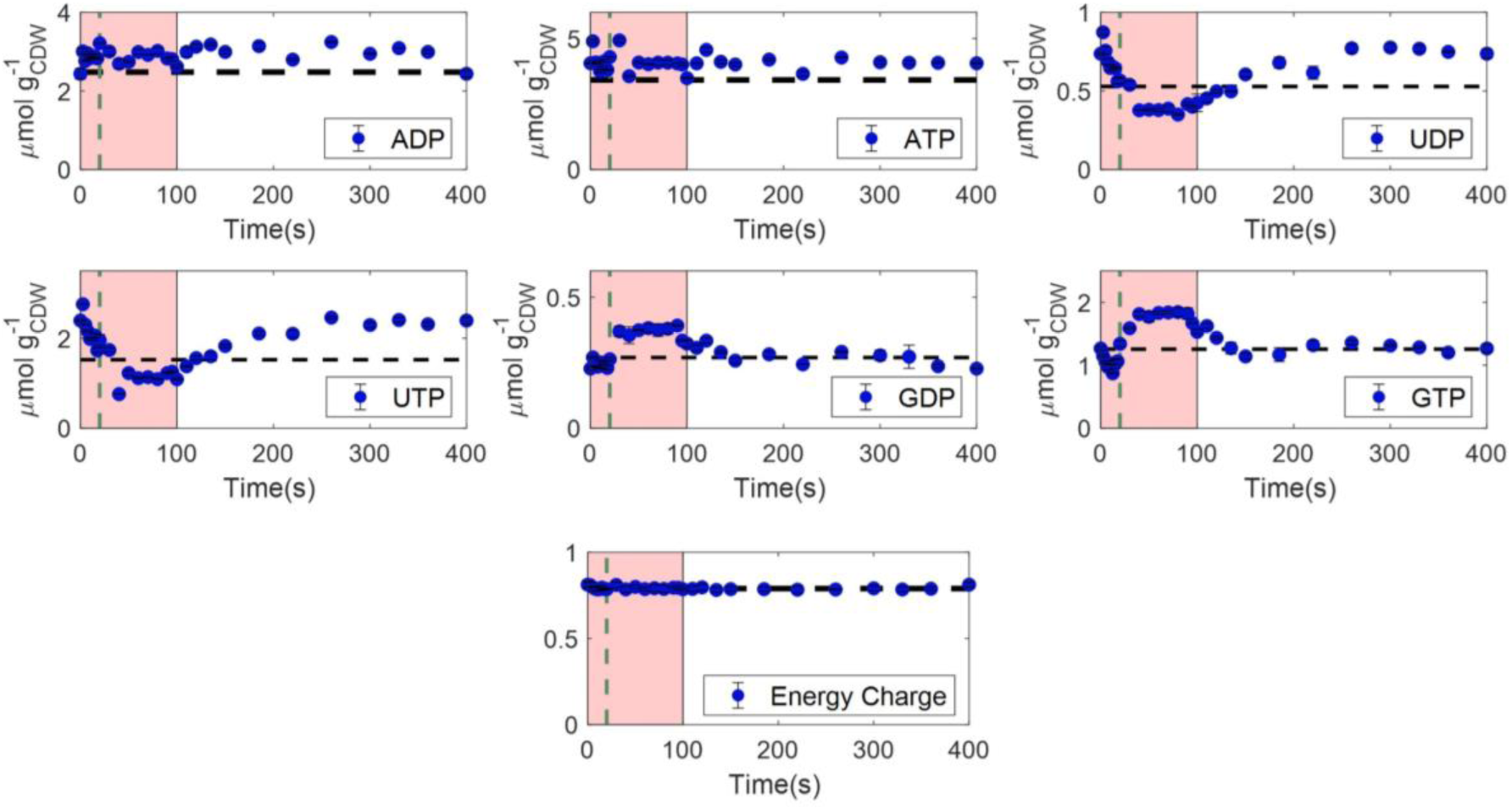
Intracellular concentrations (μmol/g_CDW_) of nucleotides, as well as the adenylate energy charge (AEC), over a feast-famine cycle (s). Black horizontal dashed lines represent the average steady-state levels. Green vertical dashed lines show the end of the feeding (20 s). The pink area represents the substrate feast phase.

#### Total Metabolome

From the extracellular observations, it was observed that the carbon uptake and excretion were significantly shifted. While the substrate carbon was consumed only during the first 100 s, excretion in the form of CO_2_ was observed over the whole cycle (400 s). The specific glucose uptake rate was higher than at the reference steady-state and there was no significant accumulation of by-products. This suggests high intracellular accumulation of carbon during the feast phase and degradation during the (extracellular) famine phase.

The total amount of intracellular carbon during steady-state and feast-famine was calculated from the measured metabolites at every timepoint and is shown in Figure 7.

**Figure 7.**
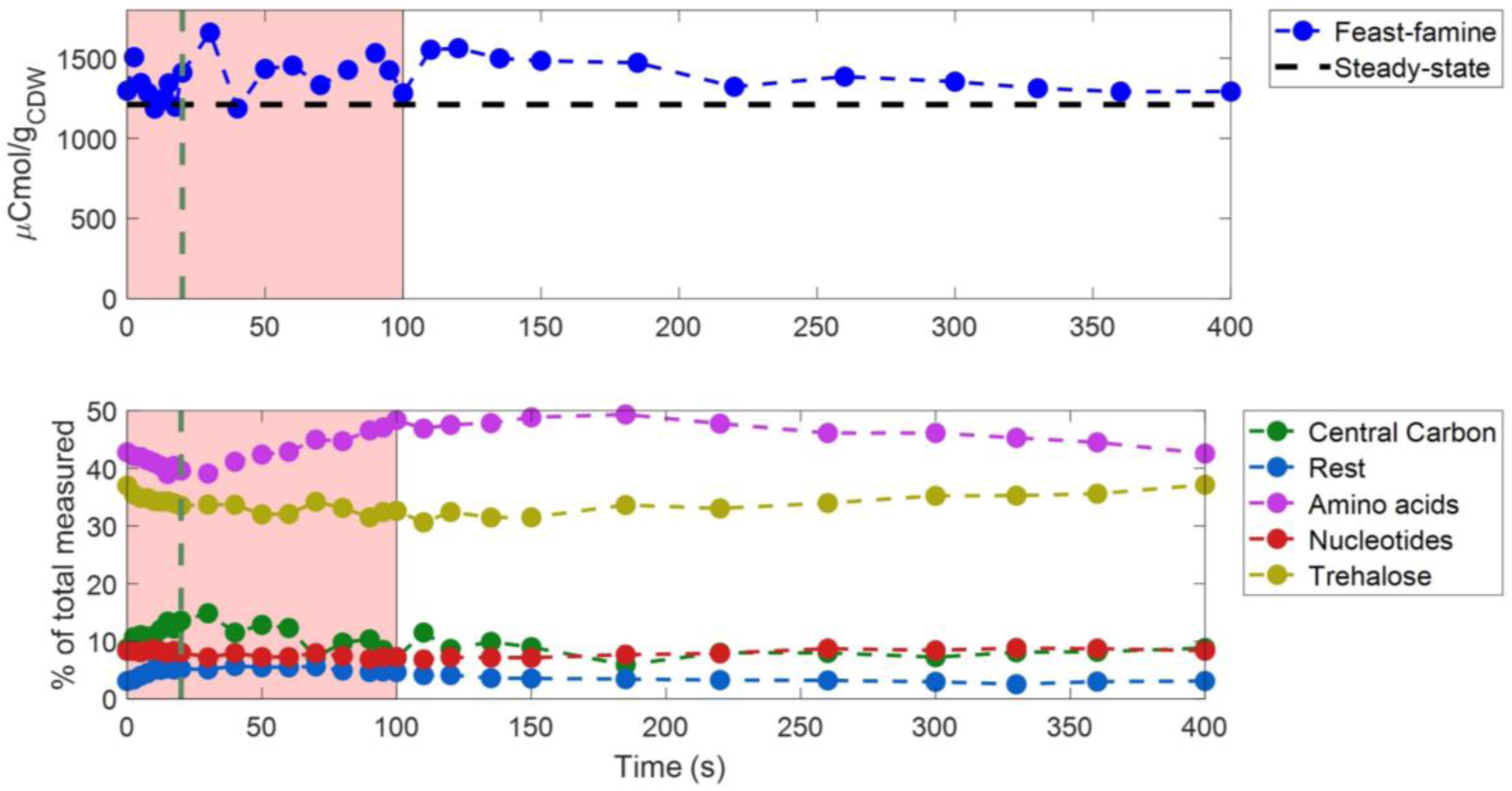
Top: Total amount of intracellular metabolites measured (in μCmol/g_CDW_) over a feast-famine cycle (s). The black horizontal dashed line represents the average steady-state levels. Bottom: The carbon distribution (% of the total intracellular metabolome measured) in metabolites of different categories/pathways, over a feast-famine cycle (s). Green vertical dashed lines show the end of the feeding (20 s). The pink area represents the substrate feast phase. The detailed list of metabolites for each category can be found in Supplementary Material S9.

The total amount of metabolites was changing over time, during the feast-famine regime. We observed a small decrease in the total metabolome during the first 10 seconds of the cycle, followed by an increase until the highest point (1661 μCmol/g_CDW_), at 30 s (Figure 7-top). After this point, the concentrations remained constant (around 1420 μCmol/g_CDW_) until approximately 120 s. Then a constant decrease, until reaching the initial level, was observed. Significantly, the total amount of carbon in the measured metabolome was, at all timepoints, higher than the steady-state levels, resulting from the overshoot in the glucose uptake.

But which metabolites accounted for the highest changes? In Figure 7 (bottom) metabolites have been divided in different categories, based on their pathways, and the carbon percentage of the total metabolome that they represented was plotted over time. Amino acids were found to contain most of the total carbon during the whole cycle, ranging from 39 to 49%, with glutamate contributing the most to this observation, being the most abundant pool measured. Trehalose was also a significant pool, accounting for approximately 34% of the total carbon, remaining however, constant over time. In the first 20 s of the feast-famine regime, the carbon percentage of central carbon intermediates (glycolysis, PPP and TCA) was increasing, in addition to the metabolites M6P, T6P, UDP-glucose, M1P and G1P (representing the rest in Figure 7-bottom). The highest change was attributed to citrate (2-6% of the total carbon measured). After 20 s, while glucose uptake rate was already lower, the filling of the amino acid pools was evident, as their carbon percentage increased during the feast, as well as the first 100 s of the famine phase. Nucleotides remained constant over time, as discussed earlier in this section.

From the oxygen uptake profile (Figure 2E), we derived that 41.5% of the total oxygen consumption occurred during the first 110 s of the cycle (feast). However, oxygen was still consumed in the famine phase, with a lower rate, until approximately 250 s when it reached the initial uptake level and remained constant until the end of the cycle. Since glucose was depleted during that phase and by-product concentrations were not changing over time, the electrons consumed must have been supplied either by an intracellular storage compound or other accumulated intermediates, as is also shown from the RQ calculation (Figure 2G).

The most common storage polysaccharides in *E.coli* are trehalose [89] and glycogen [90]. As discussed above, trehalose was a big intracellular pool, but did not change over time and was therefore ruled out as a buffer compound. Glycogen is the most well-known storage polysaccharide in *E.coli*. When the substrate is in excess, some of the glycolytic flux is diverted in the production of glycogen by G1P. The cells can then use this storage to grow under substrate limitation [91]. An attempt to quantify intracellular glycogen was performed, leading only to the conclusion that glycogen levels were increased during feast-famine, compared to steady-state. However, the measurements were not accurate enough to conclude if there was production and consumption during the dynamic cycle and therefore data are not shown.

Looking at the rest of the intracellular intermediates, the total accumulation, in terms of carbon, during the feast phase was calculated to be 256 μCmol/g_CDW_, which is 1.6 mCmol/L_EC_ (Figure 7-top). This amount represented 34% of the total glucose (4.7 mCmol/L_EC_) consumed by the cells in the first 100 s of the feast-famine cycle. In terms of electrons, this accumulation in the feast phase was 6.4 mEmol/L_EC_. If all the electrons were used during the famine phase, the maximum oxygen consumption observed would be 1.6 mmol_O2_/L_EC_. In fact, we estimated, from the calculated q_O2_, that indeed around 1.6 mmol_O2_/L_EC_ were consumed during the famine phase. Therefore, all of the accumulated intracellular metabolites could have been used as electron donors and could explain the O_2_ uptake, while the substrate was depleted. Taymaz-Nikerel H*, et al.* [52] reached to similar conclusions. In their case, 50% of the intracellular metabolites were catabolised in the famine phase, after a single glucose pulse, with glutamate being the most abundant pool.

## 4. Discussion

*E. coli* cultured under dynamic substrate conditions exhibited a different physiology compared to conditions supplying the same amount of substrate steadily. Namely, the average biomass specific consumption (glucose, O_2_) and production rates (CO_2_) increased under the feast-famine regime, compared to the steady-state, while by-product synthesis remained unaffected. Consequently, biomass formation was adversely disturbed with 30% decrease in yield.

These observations suggest that during the intermittent feeding, more glucose was used for respiration than biomass production. This would mean that either the excess of ATP produced was used in other cellular processes or the activity of the proton-translocating ATPase showed a reduced efficiency, leading to a decrease of ATP synthesis. If the energy-spilling scenario is correct, it would imply an increase in maintenance or the presence of futile cycles.

Alternatively, this change in physiology could be attributed to an increase in competitiveness. Fast consumption of the substrate generates an advantage compared to slow competitors, as an increasing share of the substrate will go to the faster consuming cells [92].

### Maintenance

Maintenance is defined as the energy-consuming processes “for functions other than production of new cell material” [93]. In *E.coli*, the most important maintenance processes described in literature, are protein turnover (synthesis and degradation), switches in metabolic pathways, proteome and RNA repair and cell motility [92, 94–96]. Any of these parameters can, therefore, provide an explanation of the decrease in biomass yield. Protein turnover rate is an important characteristic of the cell. Energetically, a majority of ATP is used for protein synthesis and degradation. It is well-known that the cell uses its proteasome to degrade misfolded proteins or proteins with other abnormalities. Therefore, the change from a steady-state to a dynamic environment may have resulted in accumulation of proteins which have to be rapidly eliminated by the cell [97]. The ATP demands for protein turnover could have thus increased, causing the loss in biomass yield. The cost of protein degradation by the proteasome has been estimated to be minimum two ATP molecules per peptide bond [98, 99]. Protein turnover rates can be quantified with dynamic ^13^C labeling of the amino acids pools and proteasome activity essays [100, 101], which were not performed in this study.

### Futile Cycles

Futile cycles may also explain the ATP-spilling during the feast-famine regime. Some potential futile cycles, already described in literature, are:

1. The reconversion of oxaloacetate to phosphoenolpyruvate by the gluconeogenic PEP carboxykinase [102]. This ATP-dissipating futile cycle has been identified for low dilution rates in glucose-limited chemostats of a pyruvate kinase deficient *E.coli* strain [103], as well as, for the wild-type under very low glucose availability [104]. The induction of this futile cycle showed stimulation of glucose and oxygen uptake rates, decrease of growth yield on glucose and increase of fermentation products [105]. Yang C*, et al.* [106] demonstrated that ATP dissipation, by the PEP carboxykinase futile cycle, increased with the decrease of the growth rate, reaching 8.2% of the total ATP produced, at a dilution rate of 0.01 h^-1^.
2. The reconversion of fructose-1,6-biphosphate to fructose-6-phosphate by fructose 1,6-bisphosphatase. Usually this futile cycle is tightly regulated by the cells and is therefore minimal under both glycolytic and gluconeogenic conditions [107, 108].
3. The reconversion of pyruvate to phosphoenolpyruvate by phosphoenolpyruvate synthase, with the involvement of Enzyme I of the PTS [109, 110]. 10% of PEP was found to be produced by pyruvate during growth of *E.coli* wild-type on glucose [109].
4. The reconversion of acetate to acetyl-coA by acetyl-coA synthetase [111]. Valgepea K*, et al.* [112] claimed that bacteria may use this futile cycle under low nutrient availability, for chemotaxis, fighting other organisms, biofilm formation and other functions.
5. Glycogen formation and re-consumption. The formation of glycogen, by glycogen synthetase, requires one ATP per glucose, while its consumption does not form any ATP [92, 113].

All of the above-mentioned futile cycles could be active in *E.coli* during the feast-famine conditions. It is well possible that the cells up-regulate the enzymes of these futile cycles, in order to rapidly switch the direction of the fluxes, when shifting from feast to famine conditions. The reasons behind this metabolic strategy can be the rapid re-initiation of growth and the tight regulation of the ATP levels in the cell. We indeed observed a constant energy cellular status over the intermittent regime, which enhances the hypothesis of this role of futile cycles. However, the identification and interpretation of these energy-spilling reactions, under alternating feast-famine conditions, has not been studied in literature. Measurements of the enzyme expression and ^13^C flux determination will be needed to confirm our hypotheses.

### Proton Translocation

If the less effective proton-translocation ATP synthase scenario is correct (reduced P/O ratio), that would mean that the cells would need to compensate for more ATP (1) by substrate-level phosphorylation and (2) increased respiration. Since the glucose uptake was higher under the feast-famine conditions, compared to the reference steady-state, more redox equivalents were produced over time, developing the need for higher respiration rates, which was also verified by our experimental data. In this case respiration is not totally coupled to ATP synthesis, as has been observed before [114]. Noda S*, et al.* [115] and Jensen PR*, et al.* [116] observed significantly decreased growth yield and increased specific rates of both glucose and oxygen consumption in different mutant *E. coli* strains, lacking F_1_-ATPase, under steady-state and batch growth, which shows how the microorganism behaves when oxidative phosphorylation is impaired. Koebmann BJ*, et al.* [117] also demonstrated similar results by manipulating the expression of the F1 subunits from the H^+^-ATP synthase, enhancing uncoupled ATP hydrolysis in *E.coli*.

However, it is still surprising that, in our experimental work, the rise in substrate-level phosphorylation did not lead to an increase in acetate formation, contrary to all the above mentioned studies. Yet, the reason behind the potential less efficient function of ATP synthase is, however, unknown. A hypothesis could be the need of the cells to translocate more protons outside of the cytosol, compared to the amount imported, in order to maintain the intracellular pH homeostasis. Due to the high metabolic rates, many acidic compounds were produced, acidifying the cytoplasmic space. Therefore, decreasing the expression of ATP synthase that brings protons into the cell, can be an advantageous strategy for the intracellular pH to remain constant and close to neutral values. This strategy has been observed for growth under highly acidic environments [118, 119]. However, the decrease in ATP synthase efficiency has not been studied for feast-famine conditions in literature, therefore a proteome study on this complex would be necessary to support this hypothesis. Another approach would be to identify the P/O ratio during steady-state and feast-famine by cultivating the cells under different substrates and growth rates as in [120–122]. More methods have also been used in literature, such as ADP pulses combined with oxygen electrode measurements (for review see [123]).

### Short-term uptake dynamics

Based on high time resolution, extracellular glucose concentration measurements and PWA rate approximations, the glucose uptake rate was calculated over cycle time and showed an immediate increase after the feeding was switched on. This rate reached a maximum value, higher than batch maximum rates, and decreased before the concentration decreased. The fact that glucose was still in excess, but the uptake rate could not follow up, indicates the existence of an intracellular metabolic limitation. From our intracellular metabolic analysis, we showed that there was enough PEP during the whole cycle to drive glucose transport in the cell through the PTS, therefore, the import of substrate was not the limiting step. One hypothesis which could explain the switch in the uptake rate is the phenomenon of macromolecular crowding. Macromolecular crowding is occurring in all living organisms, as a large part of the volume of the cell is occupied by high concentrations of macromolecules, such as proteins. Thus, there is limited intracellular volume available for other molecules. This volume exclusion affects various enzymatic reactions, either by increasing or decreasing their rates, depending on the change in size of the reactants [124–127]. While it has been shown that macromolecular crowding demonstrates mostly advantageous effects on cell metabolism [124, 127–129], it can also function as a constraint for cells which exhibit high metabolic rates [130]. Beg QK*, et al.* [131] developed a flux balance model of *E.coli* (FBAwMC), including a constraint for the enzyme concentrations, considering the macromolecular crowding. With this model they predicted the maximum growth both in single-substrate, but also in mixed substrate media, accurately representing experimental observations, showing that the growth rate was influenced by the solvent availability in the cytoplasm. In a following study, Vazquez A*, et al.* [132] applied the same modelling framework to changes from low to high growth rates. Their results demonstrated, among others, that under high metabolic rates the limitation in substrate uptake and growth rate is highly related to the crowding of the intracellular space. Therefore, the high metabolic rates observed in our study, during the feast-famine regime, may have caused the limitation observed in the glucose uptake rate after 18 s, as the cytosol space may have been unable to handle further increase of macromolecules produced. It was, however, challenging to identify if this source of regulation is indeed the cause for the change in the uptake rate, with the current dataset. More reasons could involve membrane integrity, as there is a minimal lipid to transporter proteins ratio [133] or enzyme kinetic constraints [130].

Furthermore, we observed the high capacity of intracellular metabolism facing the substrate gradients applied. There was a rapid response in intracellular metabolite concentrations and fluxes, which generally deviated significantly from the steady-state. The dynamics observed were less pronounced moving downstream, from glycolysis to TCA and then to amino acid synthesis, with more modest changes over time. One of our most interesting observations, was the impressive capability of the cells to maintain the adenylate energy charge homeostasis, over the whole time of the substrate perturbations. One potential mechanism, which could explain this balance between ATP production and consumption, is the production of inorganic polyphosphate, a long-chain polymer, as an energy buffer [134]. The enzyme polyphosphate kinase (PPK) has been identified and characterized in *E.coli* [135–137]. This enzyme is responsible for the polymerization of the terminal phosphate of ATP towards polyphosphate (nATP ↔ nADP + polyP_n_). It also catalyses the reversible reaction of ADP phosphorylation [138]. Many functions of polyphosphate have been described in literature, including ATP substitute, energy recycling, and environmental stress regulation [139–142]. It is therefore highly possible that the cells can balance the ATP production and consumption during the feast-famine regime, by synthesizing inorganic polyphosphate when ATP is produced in excess and consume it when the demand for ATP is increasing. Proteomics or enzymatic assays are necessary steps to prove the existence of polyphosphate kinase and the potential polyphosphate accumulation under these dynamic conditions.

In addition, during the highly dynamic conditions, applied in this study, we demonstrated the ability of the cells to store an amount of carbon and electrons intracellularly during the feast phase, which were then used when substrate was depleted, therefore, explaining oxygen consumption during the famine phase. This strategy proves to be important for the survival and robustness of *E.coli* under nutrient-limited conditions.

### Industrial relevance

These observations are highly relevant in an industrial context, where *E.coli* is aerobically cultivated in large-scale bioreactors, facing long-term substrate gradients. Fed-batch regimes are often preferred, since the substrate concentration or the specific growth rate can be controlled in such a way to avoid overflow metabolism [143, 144]. Therefore fed-batch cultivations facilitate higher biomass and product yields than batch or chemostat cultivations [145]. However, with the present study, we have shown that the circulation of cells around zones of substrate excess and limitation can lead to significant biomass yield losses, decreasing the profitability of the process. In addition, the immediate response of the microorganism to the excess of substrate, observed by the increased capacity of uptake rate, also leads to higher oxygen consumption. Therefore, oxygen limitation will be observed in these zones, or more oxygen should be supplemented in the process, which is not economically favourable. Moreover, several modelling approaches, which have been used to predict the behaviour of the cells in the scale-up, assume biomass yield as the optimization target. However, yields from steady-state cannot be transferred to dynamic conditions. Additional energy is required for processes like maintaining the energy charge homeostasis. Also, if these dynamic conditions affect severely the endogenous pathways of a wild-type strain, we would expect that artificial metabolic pathways would be even more sensitive, as their regulation in an engineered strain has not evolved over various environmental conditions [146].

## 5. Conclusions

Studying the physiological and metabolic responses of an adapted *Escherichia coli* culture, highlights parameters to take into account for metabolic engineering and process design in relation to large-scale reactor operation.

1. Cells responded immediately to an excess of substrate, by increasing their uptake rate and consequently the intracellular fluxes in tens of seconds. Carbon was stored in intracellular intermediates, during substrate feast and was consumed during a famine phase.
2. Despite, the highly changing dynamics, energy charge homeostasis was observed, as a remarkable fitness characteristic of the response to perturbations, indicating rapid metabolic regulation.
3. More important and highly relevant to industrial fermentations, was the 30% decrease of the biomass yield, occurring during the intermittent feeding, compared to a reference steady-state. Energy-spilling, was therefore, a trade-off for the adaptation of the microorganism in the dynamic environment, seeking for robust growth.

The obtained results revealed some reasons for the reduced performance of cell factories during scale-up. *E.coli* responds to stress, induced by substrate gradients, by launching a specific metabolic strategy. In order to improve productivity cost-effectively in large-scale bioprocesses, we need to further identify the mechanisms behind stress adaptation, limitations in substrate uptake and respiration, potential energy-spilling pathways and optimal growth targets of the cells, combining multi-omics approaches.

## Author contributions

Eleni Vasilakou performed all the experiments, processed the data and wrote the manuscript. Mark C.M. van Loosdrecht and Aljoscha S. Wahl supervised the work. All authors reviewed and approved the manuscript.

## Acknowledgements

The authors express their gratitude to Cor Ras (LC-MSMS), Patricia van Dam (GC-MS/MS) and Carol de Ram (GC-MS) for the exceptional analytical work and Johan Knoll for the TOC measurements. This research was part of the ERA-IB funded consortium DYNAMICS (ERA-IB-14-081, NWO 053.80.724).

