## Supplementary Material for "*Escherichia coli* metabolism under short-term repetitive substrate dynamics: Adaptation and trade-offs"

#### S1. Dissolved oxygen

The dissolved oxygen sensor used in this work was a polarographic ADI probe (Applisens, Applikon, Delft, The Netherlands) submerged into the broth. At a polarographic electrode, oxygen is reduced to water (cathode) and the electrons produced generate current, which transmits the signal. These types of probes are known to show some response delays, as a result of many factors, such as the membrane thickness etc. [1]. Because of the short-term behaviour of our experiment (seconds), the time delay of the probe should be taken into account, in order to estimate the real respiration rates. We will describe the oxygen probe dynamics with the following first order model [2]:

$$\frac{dC_{O2,L}}{dt} = \frac{\left(\widehat{C_{O_2,L}} - C_{O_2,L}\right)}{\tau_{probe}} \quad \textbf{(1)}$$

where $C_{O_2,L}$ is the dissolved oxygen measured by the sensor (%), $\widehat{C_{O_2,L}}$ is the estimated real dissolved oxygen in the broth (%), $t$ is the cycle time (s) and $\tau_{probe}$ is the time (s) needed for the sensor to reach 63.7 % of the ultimate response in a step exchange experiment [3]. The $\tau_{\mathrm{probe}}$ of our sensor was measured to be 16.65 s. Therefore, the estimated dissolved oxygen in the broth during the feast-famine regime was calculated as follows:

$$\widehat{C_{O_2,L}} = C_{O_2,L} + \tau_{probe}\frac{dC_{O_2,L}}{dt} \quad \textbf{(2)}$$

#### S2. Calculation of $O_2$ uptake and $CO_2$ production rates

In order to calculate the $O_2$ uptake and $CO_2$ production rates over one cycle time, the rates were first estimated by applying the respective mass balances over time.

The offgas in our system consisted of oxygen, carbon dioxide and nitrogen. Nitrogen gas was not produced or consumed during the cultivation. Therefore the sum of fractions of gases entering and exiting the reactor was 1:

$$y_{N_2,G,in} + y_{O_2,G,in} + y_{CO_2,G,in} = 1 \quad \textbf{(3)}$$

$$y_{N_2,G,out} + y_{O_2,G,out} + y_{CO_2,G,out} = 1 \quad \textbf{(4)}$$

where $y_{x,G,in}$ and $y_{x,G,out}$ are the fractions of the respective x gases ($N_2$, $O_2$ and $CO_2$) entering and exiting the reactor, respectively. The fractions of $O_2$ and $CO_2$ were measured by the offgas analyzer every minute and values for every second were obtained with interpolation.

Applying the nitrogen gas balance:

$$F_{G,in} \cdot y_{N_2,G,in} = F_{G,out} \cdot y_{N_2,G,out} \quad \textbf{(5)}$$

where $F_{G,out}$ and $F_{G,in}$ are the flow rates ($mmol_{air}\ h^{-1}$) of air exiting and entering the reactor, respectively. In our experimental setup air was provided with a flow rate of 1.875 $mmol_{air}\ h^{-1}$.

From (3), (4) and (5), the gas outflow leaving the reactor was calculated, every second of the cycle:

$$F_{G,out} = \frac{F_{G,in} \cdot \left(1 - y_{O_2,G,in} - y_{CO_2,G,in}\right)}{1 - y_{O_2,G,out} - y_{CO_2,G,out}} \quad \textbf{(6)}$$

From the mass balances of $O_2$ and $CO_2$, the rates of consumption and production were then estimated respectively:

$$R_{O_2} = F_{G,out} \cdot y_{O_2,G,out} - F_{G,in} \cdot y_{O_2,G,in} \quad \textbf{(7)}$$

$$R_{CO_2} = F_{G,out} \cdot y_{CO_2,G,out} - F_{G,in} \cdot y_{CO_2,G,in} \quad \textbf{(8)}$$

where $R_{O_2}$ ($mmol_{O2}$ $h^{-1}$) and $R_{CO_2}$ ($mmol_{CO2}$ $h^{-1}$) are the $O_2$ consumption and $CO_2$ production rates, respectively, for every timepoint in the feast-famine cycle.

The biomass specific rates were then calculated:

$$q_{O_2} = \frac{R_{O_2}}{C_{BM} \cdot V}, q_{CO_2} = \frac{R_{CO_2}}{C_{BM} \cdot V} \quad \textbf{(9)}$$

where $C_{BM}$ is the biomass concentration in the broth ($g_{CDW}$ $L^{-1}$) and V is the broth volume (L).

We performed the above calculations for 16 successive feast-famine cycles and then used the average of all cycles for every second of the cycle.

We then added a pure time delay in both rates, which was assumed to be 46 seconds for $O_2$ and 72 seconds for $CO_2$, based on the time it took for the offgas $O_2$ concentration to decrease and $CO_2$ concentration to increase (offgas analyzer) after the beginning of the feeding.

For both rates, a piecewise affine (PWA) rate approximation [4] was calculated. The breakpoints used were timepoints of 0, 20, 50, 80, 135, 262 and 400 s. These breakpoints were chosen, as they exhibited the highest goodness of fit ($R^2$ was used), among various combinations [5]. The rates between the breakpoints followed a first order linear function.

Using the measured $y_{O_2,G,out}$ and $y_{CO_2,G,out}$ ratios and the calculated $q_{O_2}$ and $q_{CO_2}$ rates, an optimization was performed (Matlab R2018a, The MathWorks, Inc.) by minimizing the sum of squares between the initial measurements and the predicted.

The following differential equations were used for the optimization:

For oxygen:

$$\frac{d[O_2]_{out}}{dt} = F_{G,in} \cdot [O_2]_{in} - F_{G,out} \cdot [O_2]_{out} - R_{O_2} \quad \textbf{(10)}$$

where $[O_2]$ is the concentration of oxygen in the gas phase.

For carbon dioxide:

At pH 7.0 there is significant interconversion of dissolved $CO_2$ and bicarbonate in the broth [6], which was taken into account in our model. Using the system described in [7], the following differential equations for $CO_2$ and $HCO_3^-$ were derived:

$$\frac{d[CO_2]_{out}}{dt} = F_{G,in} \cdot [CO_2]_{in} - F_{G,out} \cdot [CO_2]_{out} + R_{CO_2} - (k_1 + k_2 \cdot 10^{pH-14}) \cdot [CO_2]_{out} + (k_{-2} + k_{-1} \cdot 10^{-pH}) \cdot [HCO_3^-] \quad \textbf{(11)}$$

$$\frac{d[HCO_3^-]}{dt} = (k_1 + k_2 \cdot 10^{pH-14}) \cdot [CO_2]_{out} - (k_{-2} + k_{-1} \cdot 10^{-pH}) \cdot [HCO_3^-] \quad \textbf{(12)}$$

where $[CO_2]$ is the concentration of $CO_2$ in the gas phase and $[HCO_3^-]$ is the concentration of bicarbonate in the broth. $k_1$, $k_{-1}$, $k_2$ and $k_{-2}$ are the reaction constants, as described in [7]. For our calculations we used values from the literature for $37^0$C, as follows:

$k_{-1} = 60$ in $s^{-1}$ [6]

$k_1 = e^{-11.582-\frac{918.9}{T}} \cdot k_{-1}$ in $M^{-1}s^{-1}$ [8], where T = 310.15 K

$k_{-2} = 107 \cdot 10^{-5}$ in $s^{-1}$ [6]

$k_2 = \frac{e^{-11.582-\frac{918.9}{T}}}{k_{wf}} \cdot k_{-2}$ in $M^{-1}s^{-1}$ , where $k_{wf} = e^{148.9802-\frac{13847.26}{T}-23.6521 \cdot \ln T}$ is the water dissociation equilibrium constant [9].

#### S3. Extracellular by-products

**Table S.1** Extracellular by-product concentration measurements in mM for steady-state (3 replicates) and feast-famine (over time).

| | Lactate | Formate | Acetate | Ethanol |
|---|---|---|---|---|
| **Steady-state** | | | | |
| Sample 1 | 4.42 | 0.33 | 1.11 | 9.27 |
| Sample 2 | 4.53 | 0.33 | 1.12 | 8.62 |
| Sample 3 | 4.97 | 0.37 | 1.17 | 7.90 |
| **Feast-famine** | | | | |
| **Timepoints (s)** | | | | |
| 0 | 1.52 | 1.66 | 2.13 | - |
| 2.5 | 1.56 | 1.61 | 1.99 | 0.59 |
| 5 | 1.54 | 1.62 | 2.01 | 2.15 |
| 7.5 | 1.56 | 1.64 | 2.00 | 1.38 |
| 10 | 1.57 | 1.52 | 2.21 | 1.13 |
| 12.5 | 1.60 | 1.76 | 2.47 | 1.85 |
| 15 | 1.54 | 1.75 | 2.39 | 6.58 |
| 17.5 | 1.75 | 1.89 | 2.45 | 3.82 |
| 20 | 1.80 | 1.94 | 2.44 | 4.21 |
| 25 | 1.93 | 2.07 | 2.40 | 3.27 |
| 30 | 1.50 | 1.74 | 2.11 | 4.99 |
| 40 | 1.51 | 1.74 | 2.45 | 3.34 |
| 50 | 1.67 | 1.77 | 2.33 | 4.74 |
| 60 | 1.66 | 1.89 | 2.46 | 2.64 |
| 70 | 1.64 | 1.87 | 2.55 | 2.60 |
| 80 | 1.81 | 1.95 | 2.37 | 2.98 |
| 90 | 1.56 | 1.63 | 2.07 | 2.52 |
| 95 | 1.64 | 1.88 | 2.40 | 4.89 |
| 100 | 1.54 | 1.63 | 2.06 | - |
| 110 | 1.50 | 1.74 | 1.93 | 3.91 |
| 120 | 1.96 | 2.14 | 2.46 | 3.73 |

| | | | | |
|---|---|---|---|---|
| 135 | 1.56 | 1.66 | 2.13 | - |
| 150 | 1.58 | 1.67 | 2.04 | 5.03 |
| 185 | 1.58 | 1.79 | 2.30 | 5.88 |
| 220 | 1.55 | 1.64 | 2.07 | 2.68 |
| 260 | 1.56 | 1.63 | 2.06 | - |
| 330 | 1.54 | 1.63 | 2.08 | - |
| 360 | 1.61 | 1.71 | 2.06 | 4.69 |
| 400 | 1.52 | 1.66 | 2.13 | - |

### S4. Biomass specific rates (raw data)

**Table S.2** Raw data of steady-state and average feast-famine biomass specific rates with their associated standard deviations.

| | **Steady-state** | | **Feast-famine (cycle average)** | |
|---|---|---|---|---|
| **Biomass concentration** (g $L^{-1}$) | 9.71 ± 0.63 | | 6.57 ± 0.15 | |
| **Biomass growth μ** (g $g_{CDW}^{-1}$ $h^{-1}$) | 0.044 ± 0.004 | | 0.048 ± 0.009 | |
| $\mathbf{q_{Glucose}}$ ($mmol_{glc}$ $g_{CDW}^{-1}$ $h^{-1}$) | -0.70 ± 0.05 | | -1.07 ± 0.03 | |
| $\mathbf{q_{O2}}$ ($mmol_{O2}$ $g_{CDW}^{-1}$ $h^{-1}$) | -2.16 ± 0.16 | | -4.22 ± 0.19 | |
| $\mathbf{q_{CO2}}$ ($mmol_{CO2}$ $g_{CDW}^{-1}$ $h^{-1}$) | 2.21 ± 0.15 | | 4.35 ± 0.12 | |
| **Respiratory Quotient** | 1.02 ± 0.10 | | 1.03 ± 0.04 | |
| | **Raw** | **Reconciled** | **Raw** | **Reconciled** |
| $\mathbf{q_{acetate}}$ ($mmol_{ace}$ $g_{CDW}^{-1}$ $h^{-1}$) | 0.005 ± 0.0004 | 0.005 ± 0.0002 | 0.016 ± 0.003 | 0.016 ± 0.003 |
| $\mathbf{q_{ethanol}}$ ($mmol_{eth}$ $g_{CDW}^{-1}$ $h^{-1}$) | 0.039 ± 0.004 | 0.041 ± 0.003 | 0.028 ± 0.012 | 0.029 ± 0.012 |
| $\mathbf{q_{formate}}$ ($mmol_{form}$ $g_{CDW}^{-1}$ $h^{-1}$) | 0.002 ± 0.0001 | 0.002 ± 0.0001 | 0.012 ± 0.003 | 0.013 ± 0.003 |
| $\mathbf{q_{lactate}}$ ($mmol_{lac}$ $g_{CDW}^{-1}$ $h^{-1}$) | 0.021 ± 0.002 | 0.022 ± 0.001 | 0.011 ± 0.002 | 0.013 ± 0.002 |
| **Biomass Yield** ($g_{CDW}$ $g_{glc}^{-1}$) | 0.32 ± 0.04 | | 0.22 ± 0.04 | |
| **Oxygen Yield** ($mmol_{O2}$ $mmol_{glc}^{-1}$) | 3.09 ± 0.31 | | 3.94 ± 0.15 | |
| **Carbon recovery** (%) | 101.8 | | 101.6 | |
| **Electron recovery** (%) | 112.3 | | 108.7 | |

#### S5. Elemental analysis

For the elemental analysis performed, carbon (C), hydrogen (H) and nitrogen (N) were quantified with a CHN-Analyzer, phosphorus (P) was quantified with UV/VIS and sulphur (S) with ion chromatography. The measurements were performed by Mikroanalytisches Laboratorium Kolbe, Oberhausen, Germany. The oxygen content was calculated, assuming that biomass was composed only by C, H, N, P, S, O and 3% metals.

**Table S.3** Elemental composition of dried biomass in steady-state and feast-famine regime. The values represent grams of elements per 100 grams of dried biomass. Standard errors were derived by duplicate samples of each regime.

| **Composition** | **C (%)** | **H (%)** | **N (%)** | **P (%)** | **S (%)** | **O (%)** |
|---|---|---|---|---|---|---|
| **Steady-state** | 48.88 ± 0.01 | 7.56 ± 0.01 | 15.23 ± 0.01 | 5.03 ± 0.02 | 1.33 ± 0.01 | 18.99 ± 0.03 |
| **Feast-famine** | 48.69 ± 0.01 | 7.48 ± 0.01 | 15.53 ± 0.01 | 5.17 ± 0.02 | 1.25 ± 0.01 | 18.89 ± 0.03 |
| **Change (%)** | -0.39 ± 0.03 | -1.06 ± 0.19 | +1.97 ± 0.09 | +2.78 ± 0.56 | -6.02 ± 1.07 | -0.53 ± 0.22 |

#### S6. Reaction list used in FBA

| | **Flux Abbreviations** | **Enzymes** | **Reactions** |
|---|---|---|---|
| 1. | PTS | Phosphotransferase system enzymes | Glucose + PEP $\rightarrow$ G6P + $PEP_{out}$ |
| 2. | G6PDH | Glucose-6-phosphate dehydrogenase | G6P $\rightarrow$ 6PG |
| 3. | PGI | Glucose-6-phosphate isomerase | G6P $\leftrightarrow$ F6P |
| 4. | PFK | Phosphofructokinase | F6P $\leftrightarrow$ FBP |
| 5. | FBA | Fructose-biphosphate aldolase | FBP $\leftrightarrow$ DHAP + GAP |
| 6. | TPI | Triose-phosphate isomerase | DHAP $\leftrightarrow$ GAP |
| 7. | GAPD/PGK | Glyceraldehyde-3-phosphate/Phosphoglycerate kinase | GAP $\leftrightarrow$ 3PG |
| 8. | PGM | Phosphoglycerate mutase | 3PG $\leftrightarrow$ 2PG |
| 9. | ENO | Enolase | 2PG $\leftrightarrow$ PEP |
| 10. | PYK | Pyruvate kinase | PEP $\rightarrow$ $PEP_{out}$ |
| 11. | GND/RPE | 6-phosphogluconate dehydrogenase/Ribulose-phosphate 3-epimerase | 6PG $\leftrightarrow$ Xyl5P |
| 12. | GND/RPI | 6-phosphogluconate dehydrogenase/Ribose-5-phosphate isomerase | 6PG $\leftrightarrow$ Rib5P |

13. TKT1 Transketolase 1 Xyl5P + Rib5P ↔ GAP + S7P
14. TKT2 Transketolase 2 Xyl5P + E4P ↔ F6P + GAP
15. TALA Transaldolase A GAP + S7P ↔ F6P + E4P

Metabolites with the subscript "out" are products outside of the balancing space. We assume that pyruvate is the product of the PYK reaction.

#### S7. Flux balance analysis – Derived fluxes

**Table S.4** FBA estimated fluxes in $\mu mol_{substrate}/g_{CDW}/s$.

| | **Fluxes (Glycolysis)** | | | | | | | | | |
|---|---|---|---|---|---|---|---|---|---|---|
| | PTS | G6PDH | PGI | PFK | FBA | TPI | GAPD/ PGK | PGM | ENO | PYK |
| **Time (s)** | | | | | | | | | | |
| 0 | 0.005 | 0.000 | 0.017 | 0.022 | 0.034 | 0.042 | 0.076 | 0.069 | 0.067 | 0.050 |
| 2 | 4.676 | 2.681 | 2.920 | 4.905 | 4.932 | 4.990 | 10.810 | 10.243 | 10.192 | 5.149 |
| 15 | 4.404 | 1.885 | 2.123 | 3.448 | 4.019 | 4.211 | 8.908 | 8.975 | 8.981 | 4.676 |
| 18 | 1.042 | 0.382 | 0.620 | 0.854 | 0.800 | 0.784 | 1.706 | 1.742 | 1.745 | 0.717 |
| 110 | 0.022 | 0.000 | 0.018 | 0.021 | 0.008 | 0.000 | 0.000 | 0.003 | 0.005 | 0.000 |
| 400 | 0.005 | 0.000 | 0.017 | 0.022 | 0.034 | 0.042 | 0.076 | 0.069 | 0.067 | 0.050 |
| | **Fluxes (Pentose phosphate pathway)** | | | | | | | | | |
| | GND/RPE | GND/RPI | TKT1 | TKT2 | TALA | | | | | |
| 0 | 0.001 | 0.0002 | 0.0002 | 0.001 | 0.001 | | | | | |
| 2 | 1.823 | 0.874 | 0.890 | 0.924 | 0.924 | | | | | |
| 15 | 1.302 | 0.658 | 0.671 | 0.674 | 0.673 | | | | | |
| 18 | 0.248 | 0.122 | 0.123 | 0.121 | 0.121 | | | | | |
| 110 | 0.006 | 0.000 | 0.000 | 0.010 | 0.000 | | | | | |
| 400 | 0.014 | 0.0002 | 0.0002 | 0.001 | 0.001 | | | | | |

### S8. Amino acids

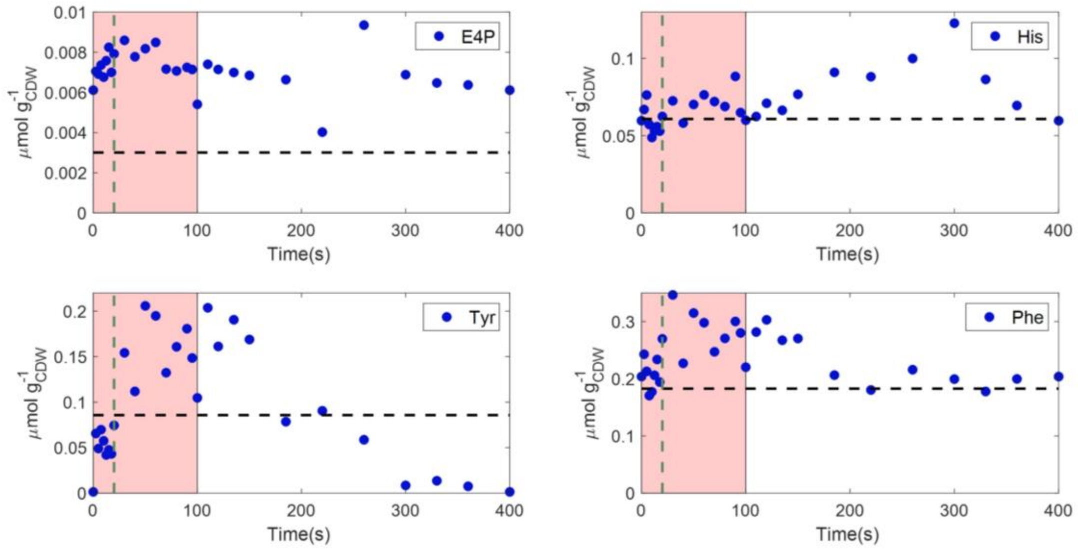


**Figure S.1** Intracellular concentrations (μmol/g$_{CDW}$) of amino acids (histidine, tyrosine and phenylalanine) with E4P as a precursor, over a feast-famine cycle (s). Black horizontal dashed lines represent the average steady-state levels. Green vertical dashed lines show the end of the feeding (20 s). The pink area represents the substrate feast phase.

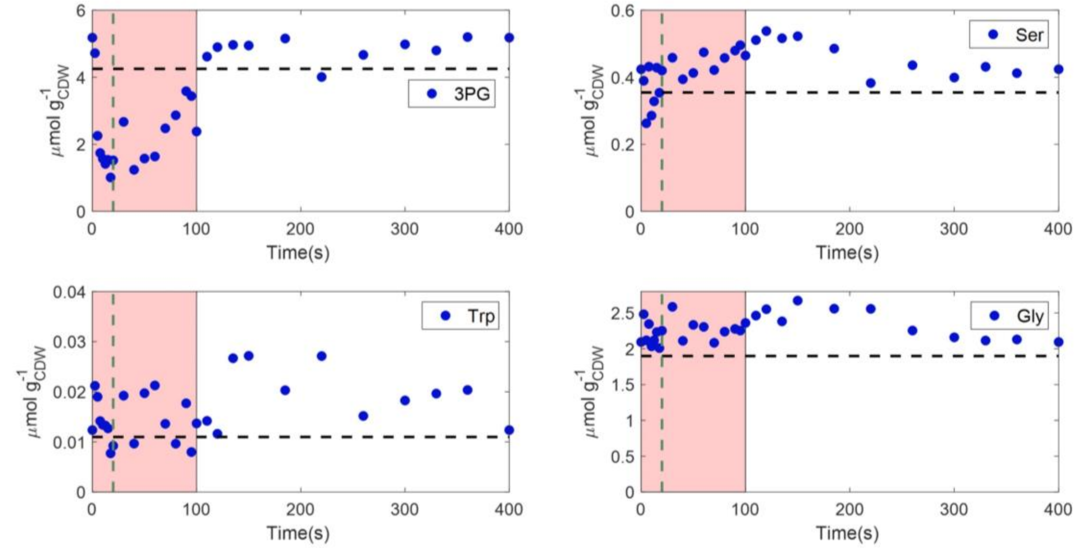

**Figure S.2** Intracellular concentrations (µmol/$g_{CDW}$) of amino acids (serine, tryptophan and glycine) with 3PG as a precursor, over a feast-famine cycle (s). Black horizontal dashed lines represent the average steady-state levels. Green vertical dashed lines show the end of the feeding (20 s). The pink area represents the substrate feast phase.

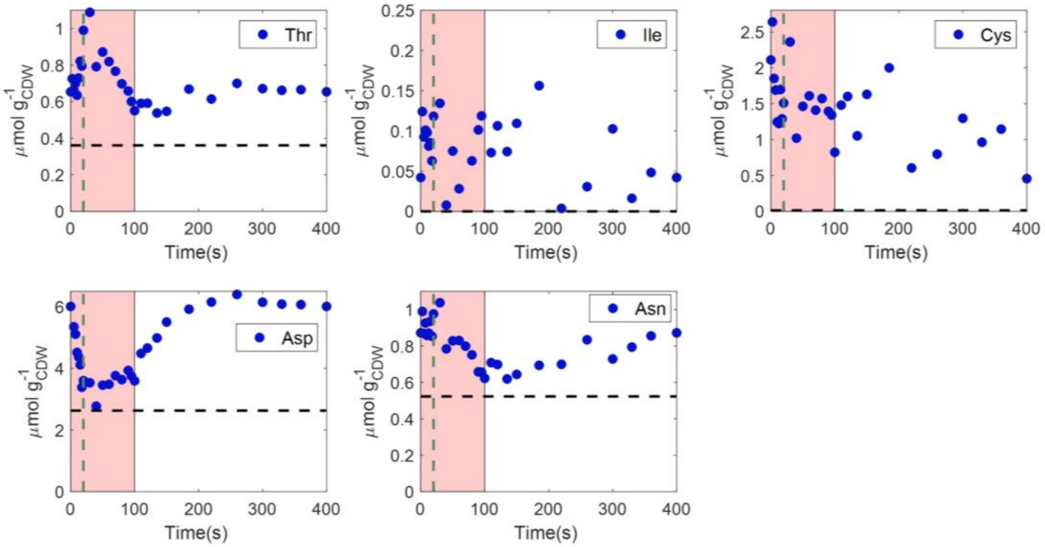


**Figure S.3** Intracellular concentrations (µmol/$g_{CDW}$) of amino acids (threonine, isoleucine, cysteine, aspartate and asparagine) with oxaloacetate as a precursor, over a feast-famine cycle (s). Black horizontal dashed lines represent the average steady-state levels. Green vertical dashed lines show the end of the feeding (20 s). The pink area represents the substrate feast phase.

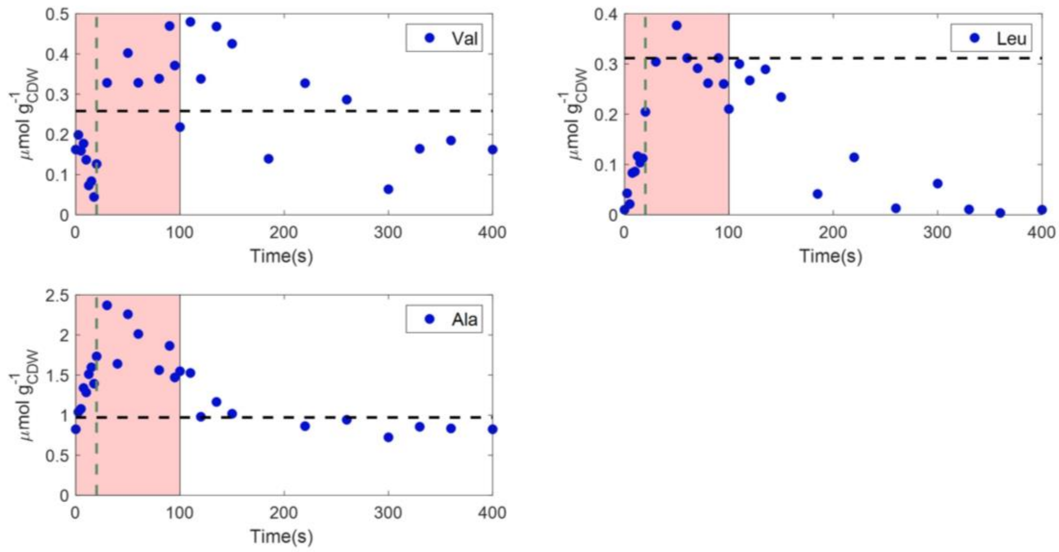


**Figure S.4** Intracellular concentrations (μmol/g$_{CDW}$) of amino acids (valine, leucine and alanine) with pyruvate as a precursor, over a feast-famine cycle (s). Black horizontal dashed lines represent the average steady-state levels. Green vertical dashed lines show the end of the feeding (20 s). The pink area represents the substrate feast phase.

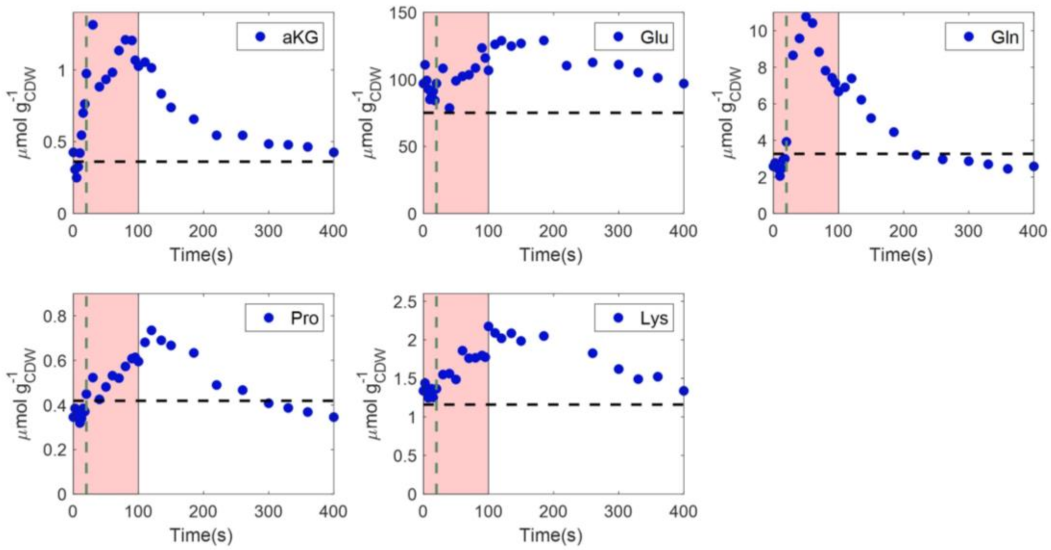


**Figure S.5** Intracellular concentrations (μmol/g$_{CDW}$) of amino acids (glutamate, glutamine, proline and lysine) with aKG as a precursor, over a feast-famine cycle (s). Black horizontal dashed lines represent the average steady-state levels. Green vertical dashed lines show the end of the feeding (20 s). The pink area represents the substrate feast phase.

#### S9. Total metabolome

**Table S.5** List of metabolites quantified in this study.

| Central Carbon | Amino acids | Nucleotides | Rest |
|---|---|---|---|
| Fumarate (Fum) | Alanine (Ala) | Adenosine diphosphate (ADP) | Trehalose (Tre) |
| Malate (Mal) | Glycine (Gly) | Adenosine triphosphate (ATP) | Trehalose-6-phosphate (T6P) |
| alpha-ketoglutarate (aKG) | Valine (Val) | Uridine triphosphate (UTP) | Mannose-6-phosphate (M6P) |
| Glyceraldehydephosphate (GAP) | Leucine (Leu) | Uridine diphosphate (UDP) | Uridine diphosphate glucose (UDP-glucose) |
| Citrate (Cit) | Isoleucine (Ile) | Guanosine diphosphate (GDP) | Mannitol-1-phosphate (M1P) |
| Isocitrate (iCit) | Proline (Pro) | Guanosine triphosphate (GTP) | Glucose-1-phosphate (G1P) |
| 2-phosphoglycerate (2PG) | Serine (Ser) | | |
| 3-phosphoglycerate (3PG) | Threonine (Thr) | | |
| Dihydroacetonephosphate (DHAP) | Methionine (Meth) | | |
| Erythrose-4-phosphate (E4P) | Aspartate (Asp) | | |
| Ribose-5-phosphate (Rib5P) | Phenylalanine (Phe) | | |
| Xylose-5-phosphate (Xyl5P) | Glutamate (Glu) | | |
| Fructose-6-phosphate (F6P) | Lysine (Lys) | | |
| Glucose-6-phosphate (G6P) | Asparagine (Asn) | | |
| Sedoheptulose-7-phosphate (S7P) | Glutamine (Gln) | | |
| Fructobiphosphate (FBP) | Tyrosine (Tyr) | | |
| Phosphoenolpyruvate (PEP) | Histidine (His) | | |
| Succinate (Suc) | Cysteine (Cys) | | |
| 6-phosphogluconate (6PG) | Tryptophan (Trp) | | |
